# Quantitative metagenomics using a portable protocol

**DOI:** 10.1101/2025.10.28.684933

**Authors:** Kaiqin Bian, Andrea Busch, John Norton, Charles Bott, Raul Gonzalez, Kyle Curtis, Dienye Tolofari, Wendell Khunjar, Katherine E. Graham, Ameet J. Pinto

## Abstract

A field-deployable DNA sequencing approach for quantitative microbial community profiling can enable rapid responses for a range of applications in the water sector – from process control to wastewater surveillance. Current quantitative approaches have complex instrumentation requirements and long turnaround time for DNA recovery and absolute quantitation. In this study, we report a field-deployable rapid detection and absolute quantitation (rD+rQ) workflow that leverages the real-time sequencing capabilities of Nanopore sequencing for quantitative metagenomics. This workflow integrates a high-molecular-weight DNA recovery protocol for diverse environmental matrices of relevance to the water sector and multiplexed Nanopore sequencing with barcoded spike-in-based calibration (BSINC). BSINC using multi-species genomic spike-in controls exhibits significantly higher calibration accuracy compared to conventional approaches that utilize either single DNA fragment or single organism spike in controls. Dynamic detection and quantitation limits were established based on coverage fraction of sequenced genomes and the coefficient of variation of genome copy numbers across replicates to enhance the accuracy and precision of microbial quantitation. The rD+rQ workflow achieves species-level identification and absolute quantitative results comparable to digital PCR in environmental samples. This portable laboratory and easy-to-use rD+rQ workflow should facilitate rapid decision-making for the water industry.

## 1. Introduction

Rapid, field-deployable and quantitative microbial profiling can enable real-time, on-site decision-making in the water sector by accelerating pathogen detection and characterizing community shifts at the point of need and thus, overcome delays inherent to centralized laboratory workflow^1,2^. Current quantitative microbial detection techniques remain constrained to laboratory-based frameworks^3,4^. Culture-based methods (e.g., heterotrophic plate counting) are time-consuming and cannot detect unculturable microorganisms^5^. Molecular techniques (e.g., quantitative polymerase chain reaction (qPCR)) require carefully designed primers and extensive parameter optimization^6^. Both organism- or gene-centric approaches cannot detect unknown or emerging organisms or capture the full complexity and dynamics of microbial communities in complex environments^6^. In contrast, shotgun DNA sequencing (i.e., metagenomics) enables comprehensive community analysis without prior knowledge of underlying community. However, the sequential workflow for metagenomics, where data analysis can only commence after the completion of the sequencing run, can introduces significant delays between sample collection and biological interpretation has constrained application for microbial detection^8,9^. These delays can undermine rapid decision-making in public health surveillance and operational control in water treatment processes^10–12^. In contrast, Oxford Nanopore Technologies sequencing technology provides a portable, real-time, and long-read alternative. Nanopore sequencing enables data analysis such as basecalling and taxonomic classification during sequencing, dramatically reducing the sample-to-answer turnaround time to 6 hours^13–18^. In addition, the long-read capability of Nanopore sequencing supports taxonomic identification at a higher phylogenetic resolution^19^. Yet, two key factors still limit the utility of Nanopore sequencing for decision making in the water sector.

First, the lack of a portable DNA extraction workflow to recover high-molecular-weight DNA (HMW DNA) from environmental samples remains the bottleneck for on-site Nanopore sequencing^20,21^. Conventional extraction methods (e.g., Qiagen DNeasy PowerWater, Qiagen DNeasy PowerSoil Pro) compatible with Nanopore sequencing are time-intensive and complex instrumentation (e.g., high-speed centrifuge, high-speed homogenizer) which limits field deployment^20–22^. Rapid in-field DNA extraction methods fail to produce DNA quantity and/or quality that is both required for and leverages the benefits of long-read sequencing^21,24–26^. Thus, to advance the applicability of long-read Nanopore sequencing on-site, it is essential to develop an efficient and field-deployable DNA extraction method that both meets the DNA requirements of Nanopore sequencing and also provides HMW DNA from environmental samples.

Second, current Nanopore-based sequencing approaches do not enable absolute quantitation but provide relative abundance information. Although relative quantitation facilitates the characterization of microbial composition, the lack of data on total microbial loads (e.g., total cells, total 16S rRNA gene concentration) limits the comparability between different studies ^28,29^. Existing methods for absolute quantitation (e.g., qPCR, flow cytometry, heterotrophic plate counting) require complex instrumentation (e.g., quantitative PCR system, flow cytometer)^27^. While single DNA spike-in-based quantitation methods have been reported, they overlook biases from GC content, fragment size, etc. between the spike-in and the microorganisms present in the samples.^5,28^. Cellular spike-in controls risk contaminating downstream analysis due to the difficulty in distinguishing them from unknown environmental samples^29^. Furthermore, accurate quantitation of microorganisms in environmental samples also requires the establishment of detection and quantitation thresholds. Conventionally, the limits of detection (LOD) and quantitation (LOQ) are established at a fixed target organism concentration, relying on the assumptions of consistent DNA recovery rates, comparable microbial diversity within similar sample types, and especially fixed sequencing efforts^30–32^. However, these assumptions do not hold when there is substantial inter-sample and inter-experiment variability in the environmental samples and flexible sequencing efforts using Nanopore sequencing^33^.

To address these challenges, we developed a portable rapid detection and absolute quantitation workflow (rD+rQ) based on multiplex Nanopore sequencing, including an in-field DNA extraction method, mapping-based taxonomic identification, and a barcoded DNA spike-in-based calibration strategy (BSINC). Dynamic approaches for determining the limits of detection and quantitation were also developed to ensure flexibility for diverse applications. We benchmarked the rD+rQ workflow against digital PCR using both mock communities and environmental samples. This field-deployable, Nanopore sequencing-based framework requires minimal infrastructure while providing high-resolution taxonomic profiling and absolute quantitation of multiple microorganisms simultaneously, which can support informed decision-making across the water sector.

## 2. Materials and Methods

### 2.1. Sample collection

Samples analyzed in this study include mixed liquor (ML) from a wastewater treatment plant (WWTP) in Michigan, secondary effluent (SE) that serves as input to a water reclamation system (WRS) in Virginia, and mix tank effluent from a microalgal cultivation (AL) system for nutrient recovery in Wisconsin, respectively. Samples with low microbial cell concentrations (i.e., SE) were concentrated to a volume ranging from 500 µL to 250 µL using the 0.05 μm PS Hollow Fiber Filter Concentrating Pipette system (InnovaPrep, MO, USA), achieving an effective concentration factor of roughly 2000-fold. All samples were shipped on ice to Georgia Institute of Technology and stored at -20 °C prior to DNA extraction.

### 2.2. DNA extraction protocols

We adapted three commercial DNA extraction kits (i.e., PureLyse, DNeasy PowerSoil Pro, and DNeasy PowerWater) for portable DNA extraction and compared their performance to the laboratory-based extraction results. To adapt these protocols for field-deployable extraction, conventional benchtop instruments such as the homogenizer and high-speed centrifuges were replaced with a handheld bead beater (i.e., SuperFastPrep-2 (MP Biomedicals LLC, CA, USA)) and a platform which contains high-speed microcentrifuge, PCR thermal cycler, and gel electrophoresis apparatus with LED transilluminator (Bento Lab (Bento Bioworks Ltd., London, United Kingdom)). All three portable DNA extraction strategies employ a combination of chemical and physical cell lysis. The physical cell lysis intensity can be controlled by the power level of the SuperFastPrep-2. We tested two different power settings (15 and 20) for the PowerSoil and PowerWater methods (see **Text S1** and **Figure S1** for additional information on the choice of power setting). Overall, five portable field-deployable DNA extraction strategies were compared to four laboratory-based DNA extraction strategies (**Table 1**)

**Table 1.** Overview of field-deployable and laboratory-based) DNA extraction strategies. (NA = not applicable)

| Protocol type | Protocol | Extraction Kit | Key equipment<sup>1</sup> | Cost (USD/extraction)<sup>2</sup> | Time single extraction, for multiplexing (mins/extraction)<sup>3</sup> |
| --- | --- | --- | --- | --- | --- |
| Portable | PL | PureLyse, | NA | ~ 13 | 10, 10 |
|  | PPS15 | DNeasy PowerSoil Pro | Handheld bead beater, Bento Lab | ~ 10 | 35, 5 |
|  | PPS20 |  |  |  |  |
|  | PPW15 | DNeasy PowerWater |  | ~ 13 | 35, 5 |
|  | PPW20 |  |  |  |  |
| Laboratory-based | PS | DNeasy PowerSoil Pro | Homogenizers, High speed centrifuge, Qiacube (optional) | ~ 10 | 35, 5 |
|  | PW | DNeasy PowerWater |  | ~ 13 | 35, 5 |
|  | PM | DNeasy Powermax Soil | Centrifuge with 50 mL adapter, refrigerator (optional) | ~ 36 | 50, 5 |
|  | ZM | ZymoBIOMICS DNA Miniprep | Homogenizers, High speed centrifuge | ~ 7 | 30, 5 |

**Notes:**
Protocol abbreviations: PL=PuraLyse, PPS15= Portable PowerSoil-Lysis power level 15, PPS20= Portable PowerSoil-Lysis power level 20, PPW15= Portable PowerWater- Lysis power level 15, PPW20= Portable PowerWater- Lysis power level 20, PS= PowerSoil Pro, PW= PowerWater, PM= PowerMax Soil, ZM= ZymoBIOMICS DNA Miniprep
1. Vortex, pipettes, tips, and tubes are not listed in this table as they are required for all methods. 2. Estimation of cost only considers reagents and consumables and may vary depending on the manufacturer's set pricing. 3. "Time for multiplexing" indicates the extra time required when processing multiple samples together. 4. Time estimation may vary for different operators.

All nine DNA extraction techniques were tested on three types of samples (i.e., ML, SE, and AL) in triplicate. Negative controls containing UltraPure DNase/RNase-Free Distilled Water (Invitrogen, CA, USA) were processed in parallel with the samples. DNA yield was quantified on the Qubit Flex fluorometer (Invitrogen, CA, USA) using the 1X Qubit dsDNA High Sensitivity Assay Kit (Invitrogen, CA, USA) following manufacturer instructions. Meanwhile, the extracted DNA was evaluated for DNA purity (A260/280 and A260/230) using NanoDrop 2000 spectrophotometer (Thermo Scientific, MA, USA) and for fragment distribution using Fragment Analyzer system with High Sensitivity Large Fragment 50 kb Kit (Agilent Technologies, CA, USA). Data from the Fragment Analyzer was analyzed and visualized using ProSize and R package, bioanalyzeR.

### 2.3. Amplicon sequencing and data processing

Select DNA extracts were submitted to Georgia Institute of Technology’s Molecular Evolution Core for amplicon sequencing on the Illumina MiSeq platform with V3 chemistry using primer sets of 515F/806R (Forward: 5′-GTGYCAGCMGCCGCGGTAA-3′, Reverse: 5′- GGACTACNVGGGTWTCTAAT-3′) and V4f/V4r (Forward: 5′- CCAGCASCYGCGGTAATTCC-3′, Reverse: 5′-ACTTTCGTTCTTGAT-3′) targeting the V4 hypervariable regions of the 16S and 18S rRNA genes, respectively^34,35^. Sequencing data was processed using DADA2 v3.14^36^. Briefly, the raw data was trimmed and dereplicated following the removal of reads with ambiguous bases (maxN = 0) and errors (maxEE = 2), as well as the truncation of reads with a quality score less than or equal to truncQ criteria (truncQ = 10 for 16S rRNA gene reads and truncQ = 2 for 18S rRNA gene). Filtered forward and reverse reads were merged to obtain the full denoised sequences. Merged reads outside the expected length range (250–256 bp for 16S rRNA gene and 250–400 bp for 18S rRNA gene) were discarded, and the remaining reads were subject to chimera removal, followed by taxonomic assignments using the SSU 138.1 and 132 reference databases for 16S and 18S rRNA genes, respectively^37,38^. Potential contaminating sequences were inferred by comparing sample reads to those in blank controls using the R package, Decontam^39^, and removed. For downstream statistical analysis, 16S and 18S rRNA gene amplicon data were rarefied to the minimum total sequence count across all samples (n = 16400, 16S rRNA gene; n = 1185, 18S rRNA gene). The alpha (Chao1, Shannon, and Simpson indices) and beta diversity (non-metric multidimensional scaling (NMDS) with Bray-Curtis distance matrix) analysis was performed using the package phyloseq v1.40.0 and permutational multivariate analysis of variance (PERMANOVA) was performed using the package adonis. All visualizations were generated using the package ggplot2^40^.

### 2.4. Multiplex Nanopore sequencing for rapid detection and absolute quantitation

We developed a barcoded Spike-in-based calibration (BSINC) strategy based on multiplex Nanopore sequencing to enable simultaneous detection and quantitation of microbial taxa. BSINC involves the ligation of unique barcodes to spike-in controls and samples separately, followed by pooling and sequencing together. ZymoBIOMICS Microbial Community DNA Standard II (Log Distribution, **Table S1**) (Zymo Research Corporation, CA, USA) (Zymo Log DNA), which is a mixture of genomic DNA of eight bacterial and two fungal strains, was utilized as the spike-in control. DNA extracted from ZymoBIOMICS Microbial Community Standard (Even Distribution) (Zymo Even Cell, **Table S2**) and Gut microbiome standard (Zymo Gut, **Table S3**) (Zymo Research Corporation, CA, USA), ML, SE, and AL using the optimized DNA extraction protocol (see section Results and Discussion 3.1) was used as the sample DNA. Zymo Even Cell and Zymo Gut include 10 and 21 microbial strains, respectively, with varying abundances to simulate complex environmental samples. All extracted DNA was purified by 1X AMPure XP beads before library preparation.

Multiplex sequencing of the spike-in controls and samples was performed on the MK1C platform (**Figure 1A)**. DNA of spike-in controls and samples was divided and ligated with three different barcodes provided in the SQK-RBK114.24 kit, respectively. The barcoded DNA from the spike-in control and samples were then pooled together and processed for clean-up, adapter ligation, priming, and library loading following the protocol of SQK-RBK114.24 for R10.4.1 Flow Cell on MK1C. Raw sequencing signals were decoded using the built-in Dorado basecaller (v0.7.0) with the fast model during sequencing. Low-quality (Q value < 7) or short (length < 200 b) reads were removed using Dorado in real-time. The remaining reads were demultiplexed and trimmed for adapters and barcodes for downstream taxonomic identification and absolute quantitation.

**Figure 1.**
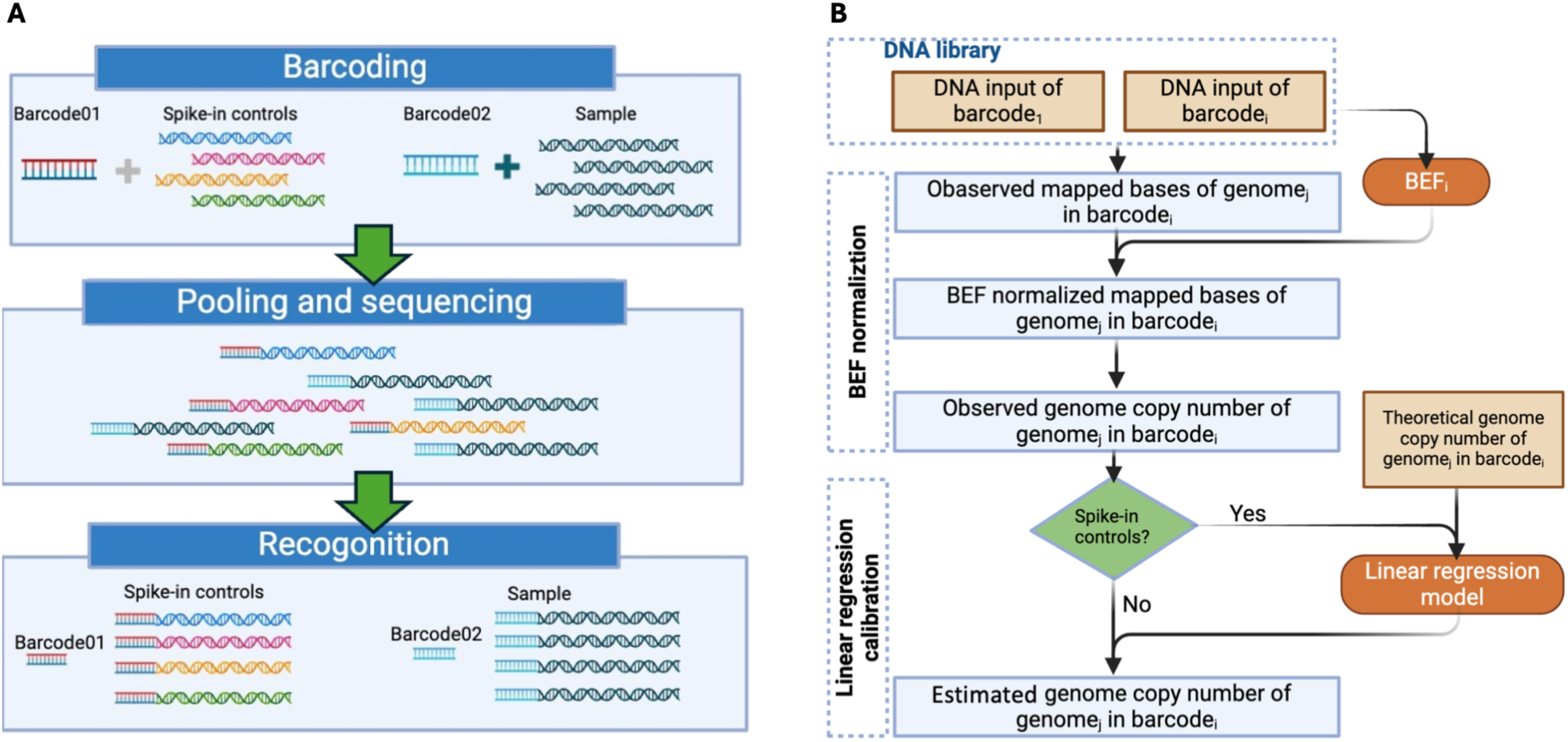
Schematic showing the overall barcode-based rD=rQ Workflow. (A) Multiplex Nanopore sequencing, detection, and (B) calibration. Spike-in controls and samples represent Zymo Mock Community DNA standard and DNA extracted from samples, respectively. BEFi indicates the barcode effect factor of barcodei.

### 2.5. Data analysis for taxonomic identification and absolute quantitation

Nanopore reads were mapped to reference databases using minimap2 (v2.24, flag: map-ont -x, mapping mode) with default parameters. For a genome to be considered detected in a sample, at least 10% of the target reference genome was required to be covered by at least one sequenced read^41^. CoverM was used to calculate the genome-wide mapped bases, covered bases, and reference genome size against the complete or draft genomes in the reference database^42^. Definitions for mapped bases, covered bases, sequencing depth, coverage fraction, and sequencing efforts are provided in **Text S2**. Reference databases for Zymo mock community (i.e., Zymo Log DNA, Zymo Even DNA, Zymo Even Cell, and Zymo Gut) were provided by Zymo. Reference databases for environmental samples (i.e., ML, SE, and AL) were built for each sample type using metagenomic sequences derived from samples of relevant water utilities (**Text S3, Figure S2**). Barcode effect factor (BEF) was first applied to each genome to normalize the batch effects for different barcodes (**Text S4, Eq. S1, S2**). BEF represents the ratio of observed sequencing effort (bp) and expected sequencing output (bp) based on input DNA (ng). The observed genome copy number for each genome, excluding barcode effects, was calculated by dividing the BEF- normalized mapped bases by the covered bases of each genome (**Text S4, Eq. S3**). To determine the absolute abundance of each genome in the samples, a linear regression model generated from spike-in controls was used to estimate the observed genome copy number of each genome. The linear regression model was derived from the correlation between the theoretical and observed genome copy numbers of the spike-in controls that are co-sequenced with samples.

### 2.6. Digital PCR

We performed digital PCR for select taxa to validate quantitative estimates obtained by the aforementioned workflow. Specifically, we targeted four commonly detected taxa at genus level (*Mycobacterium spp.*, *Accumulibacter spp.*, *Nitrospira spp., Gordonia spp.*) in ML and SE samples^43–46^. The rD+rQ workflow and dPCR measurement were conducted on the same samples simultaneously. Two sets of gBlock standards (Integrated DNA Technology gBlocks Gene Fragments) targeting the 16S rRNA gene of the target taxa were used as positive controls. The primers, probe, gBlocks, and cycling conditions were listed in **Table S4**. Technical triplicates were run (three wells) with positive (gBlocks, **Table S4**) and negative controls (UltraPure Water) on QIAcuity Digital PCR System (Qiagen, Hilden, Germany). The dPCR assays were prepared according to the manufacturer’s protocol. Triplicate wells were merged for data analysis. For each taxon, common thresholds (as listed in **Table S4**) were manually set at mean fluorescence intensity values at the peak counts of the positive and negative populations.

## 3. Results and Discussion

### 3.1. A portable DNA extraction strategy was validated across varying microbial loads and sample types

We tested the DNA extraction efficiency of five portable DNA extraction strategies (**Table 1**, PL, PPS15, PPS20, PPW15, and PPW20) across three different sample types (ML, SE, and AL) representing a range of microbial loads, water matrices, and microbial compositions. Mechanical lysis methods were key for portable DNA extraction to reduce DNA extraction time, while being supplemented with compatible chemical reagents to enhance DNA yield and microbial diversity recovery^43,47,48^. In parallel, four laboratory-based DNA extraction strategies (PS, PW, PM, and ZM) were also performed using the same samples to compare against the in-field methods. Four of the five portable DNA extraction methods (PPS15, PPS20, PPW15, and PPW20) yielded DNA in excess of 200 ng sequencing requirement for all water matrices (ML, AL, and SE) (**Figure 2A, S3A, S4A**) which exceeds the minimum requirement for rapid barcoding sequencing on a Flongle Flow Cell or standard Flow Cell (200 ng per barcode). DNA yields from 0.3 mL ML samples showed no significant difference (*p* = 0.46, Mann-Whitney U test) between four portable extraction methods (PPS15, PPS20, PPW15, PPW20: 1743.33 ± 270.50 to 3208.33 ± 67.99 ng, respectively) and three laboratory-based methods (PS, PW, PM: 2021.67 ± 86.63 to 2332 ± 362.76 ng, respectively). Comparable DNA yields between portable and laboratory-based methods were also observed in SE (*p* = 0.21, Mann-Whitney U test) and AL samples (*p* = 0.37, Mann-Whitney U test). In addition, PPS15, PPS20, PPW15, and PPW20 yielded DNA with average A260/280 ratios of 1.80−1.94, which meets the quality requirement of Nanopore sequencing. The other portable method, PL, yielded significantly lower DNA quantities (ML: 217.33 ± 33.61 ng, SE: 58.33 ± 11.51 ng, and AL: 116.27 ± 14.40 ng) than PPW15, representing only 11.76–23.76% of PPW15 DNA yields. The DNA yield of PL is constrained by the binding capacity of its beads, as it is primarily designed for amplification-based applications. While PL offers advantages in portability and processing time (**Table 1**), its DNA yield is insufficient for downstream Nanopore sequencing. These findings demonstrate that PPS15, PPS20, PPW15, and PPW20 are as effective as laboratory- based methods for on-site DNA recovery.

**Figure 2.**
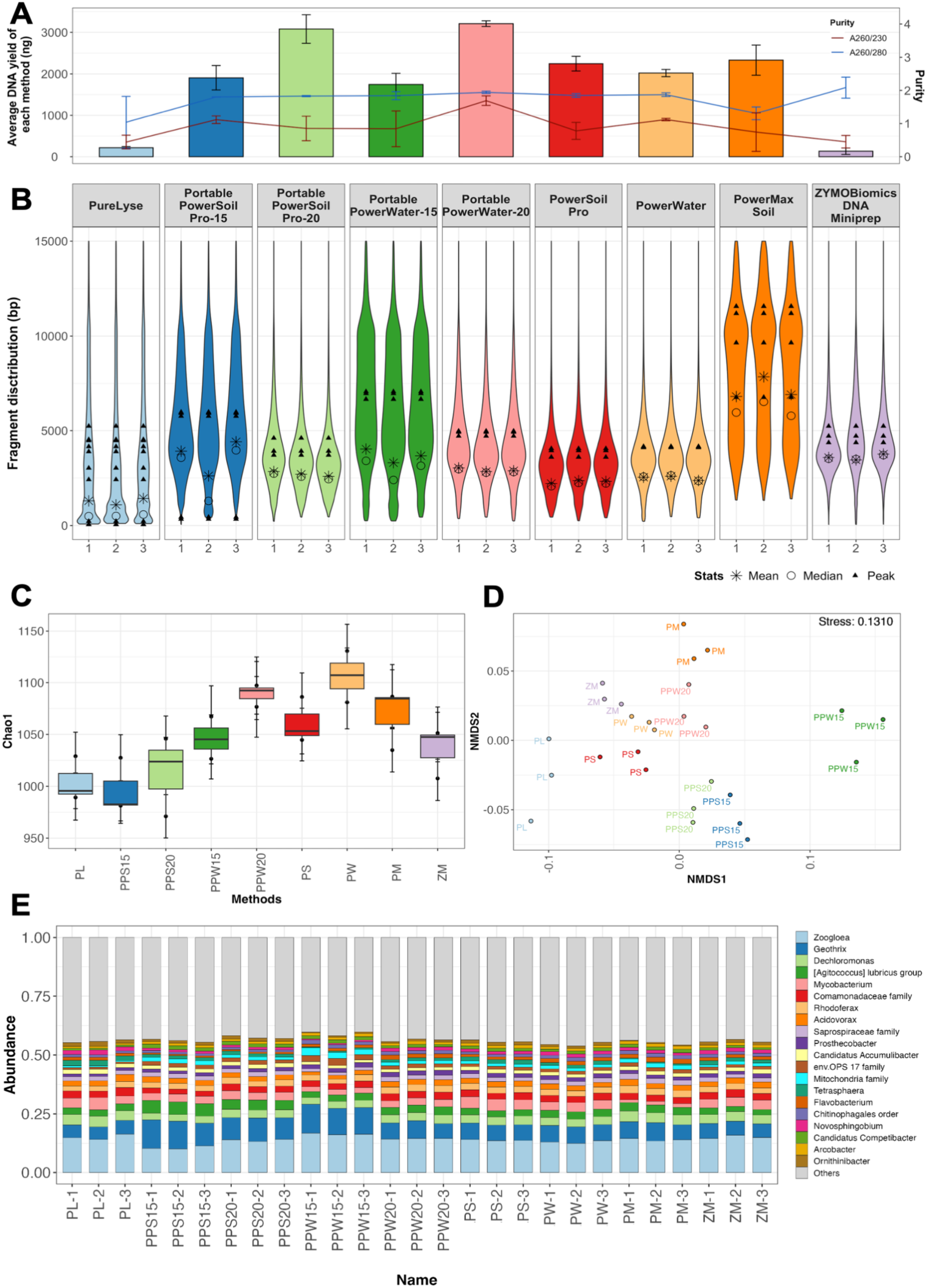
Evaluation of DNA extraction methods on the microbial community in mixed liquor sludge samples (ML). (A) Purity and quantity, and (B) DNA fragment size distribution for DNA extracted using different protocols. The line, bar, and bubble plots indicate the average purity, DNA yield, and fragment distribution, respectively. The star, circle, and filled triangle up represent the mean, median, and peak of DNA fragments by molarity, respectively. (C) Alpha diversity (Chao1 index), (D) Beta diversity, and (E) composition of the microbial community for prokaryotes. The Beta diversity is measured using NMDS2 plot based on Bray-Curtis distance.

While the DNA yields of the PPS15, PPS20, PPW15, and PPW20 met the input DNA mass for Nanopore sequencing, the DNA fragment distribution between them was highly variable. PPW15 extracted DNA demonstrated higher peak DNA fragment lengths (6895.33 ± 210.81 bp) compared with other in-field methods (58–5974 bp) and three laboratory-based methods (3606−6760 bp) from ML samples (**Figure 2B)**. Similarly, longer peak DNA fragment lengths were also observed in DNA extracted using PPW15 from AL and SE samples (**Figures S3B** and **S4B**). The results suggest that PPW15 can lyse cells without over-fragmentation of the DNA strands. PM, which utilizes gentler lysis based on vortexing, achieved the highest fragment size among all nine methods. However, it necessitates a non-portable 50 mL tube-compatible centrifuge and an additional concentration step prior to Nanopore sequencing due to its large working volume (50 mL) and dilute DNA concentration from extensive elution volume (2 mL) compared to PPW15 (2 mL and 50 μL). In addition, vortex-based lysis is more time-consuming (10 minutes) compared to the SuperFastPrep-2-based lysis (2 minutes). Therefore, the PPW15 was considered the most optimal method among all in-field methods for on-site application due to a combination of high DNA yields, high purity DNA, and larger peak DNA fragment lengths, which are critical for subsequent data analysis^47^.

16S and 18S rRNA gene amplicon sequencing, targeting prokaryotic and eukaryotic communities, respectively, was conducted to characterize the effect of extraction protocols on microbial community structure and composition. 18S rRNA gene sequencing data generated from DNA extracts of ML and SE samples using PureLyse (i.e., MLPL and SEPL) were excluded due to insufficient read numbers. Differences in alpha-diversity (Chao 1 index) were observed when comparing the same community and sample type obtained by different extraction methods (Kruskal-Wallis rank sum test, *p* = 0.034) (**Figures 2C, S3C, S4C, S5A, S6A,** and **S7A**). However, post-hoc pairwise Wilcoxon tests did not identify any significant differences between specific pairs of methods (p > 0.05). Beta-diversity analysis revealed no significant differences in prokaryotic or eukaryotic community across extraction methods for each water matrix (PERMANOVA, *p* > 0.05) (**Figures 2D, S3D, S4D, S5B, S6B,** and **S7B**). The top 20 abundant taxa were consistently detected across all extraction methods (**Figures 2E, S3E, S4E, S5C, S6C,** and **S7C**). DESeq2 results also confirmed no significant change in the relative abundance of abundant ASVs (> 1%) between different extraction methods. Previous research has primarily examined the impacts of DNA extraction methods on microbial community profiles and underlying mechanisms within the single domain (e.g., due to the different cell wall thickness and various abilities to form spores)^20,46,47,49,50^. While DNA extraction methods are known sources of variations in microbial community characterization, with eukaryotes being more susceptible to extraction biases due to their diverse cell envelopes, PPW15 captured microbial community composition similar to that of laboratory-based extraction methods for both prokaryotic and eukaryotic microbial communities in the tested sample matrices^49^.

Considering that PPW15 also results in sufficient DNA yield and the provides the longest fragment size distribution among all tested portable methods across different water matrices, it was selected as the on-site DNA extraction method for Nanopore sequencing. Unlike laboratory-based DNA extraction, which allows for standardized preprocessing to normalize biomass concentrations and remove contaminants, or customized protocols for different sample types, on-site DNA extraction in the water sector has to contend with variable biomass loads and water matrix effects, and heterogeneous microbial compositions. While the vortex mixer is portable and enables on-site mechanical bead-beating, vortexing-based lysis does not achieve consistent extraction efficiency across all sample types, especially in matrices with extremely low biomass or high levels of inhibitory substances. PPW15 addresses these challenges through a single protocol capable of extracting DNA from prokaryotic and eukaryotic communities across diverse environmental samples^51,52^. Furthermore, while mechanical lysis may compromise fragment length, we systematically identify optimal bead-beating parameters that minimize fragmentation while maintaining DNA quantity and microbial community composition recovery. Compared to methods from previous research using fixed mechanical forces without optimization (e.g., PW, PS), PPW15 achieves superior extraction efficiency^8,21,51,53^.

### 3.2. Validation of Barcoded Spike-in-based Calibration (BSINC) strategy for taxonomic identification and absolute quantitation

Spike-in controls (Zymo Log DNA) and the DNA extracted from Zymo Gut (ZG, treated as samples in this validation experiment) were ligated with three different barcodes and pooled into a single multiplexed library for Nanopore sequencing. In total, 4.78 Gbp of data were generated with an N50 of 3801 bp and a mean read quality of Q 8.9. All expected genomes of spike-in controls and ZG sample were identified in the dataset.

To validate the BSINC strategy for absolute quantitation, we compared the linear regression models between theoretical and observed genome copy numbers for spike-in controls and ZG sample across different sequencing efforts by subsampling the sequencing dataset from 10 to 10,000 Mbp (**Figure 3A**). As the total sequencing effort increased, the slope and intercept of regression models of the spike-in controls and Zymo Gut converged, demonstrating that the linear regression model derived from spike-in controls provides valid quantitative parameters for samples provided sequencing effort is appropriate. However, while the R^2^ of models for spike-in controls was greater than 0.9, these were consistently lower than 0.6 for ZG sample irrespective of the sequencing effort. This is likely due to the presence of several low-abundance genomes in the ZG sample (**Tabe S3**) relative to the spike-in control (**Table S1**).

**Figure 3.**
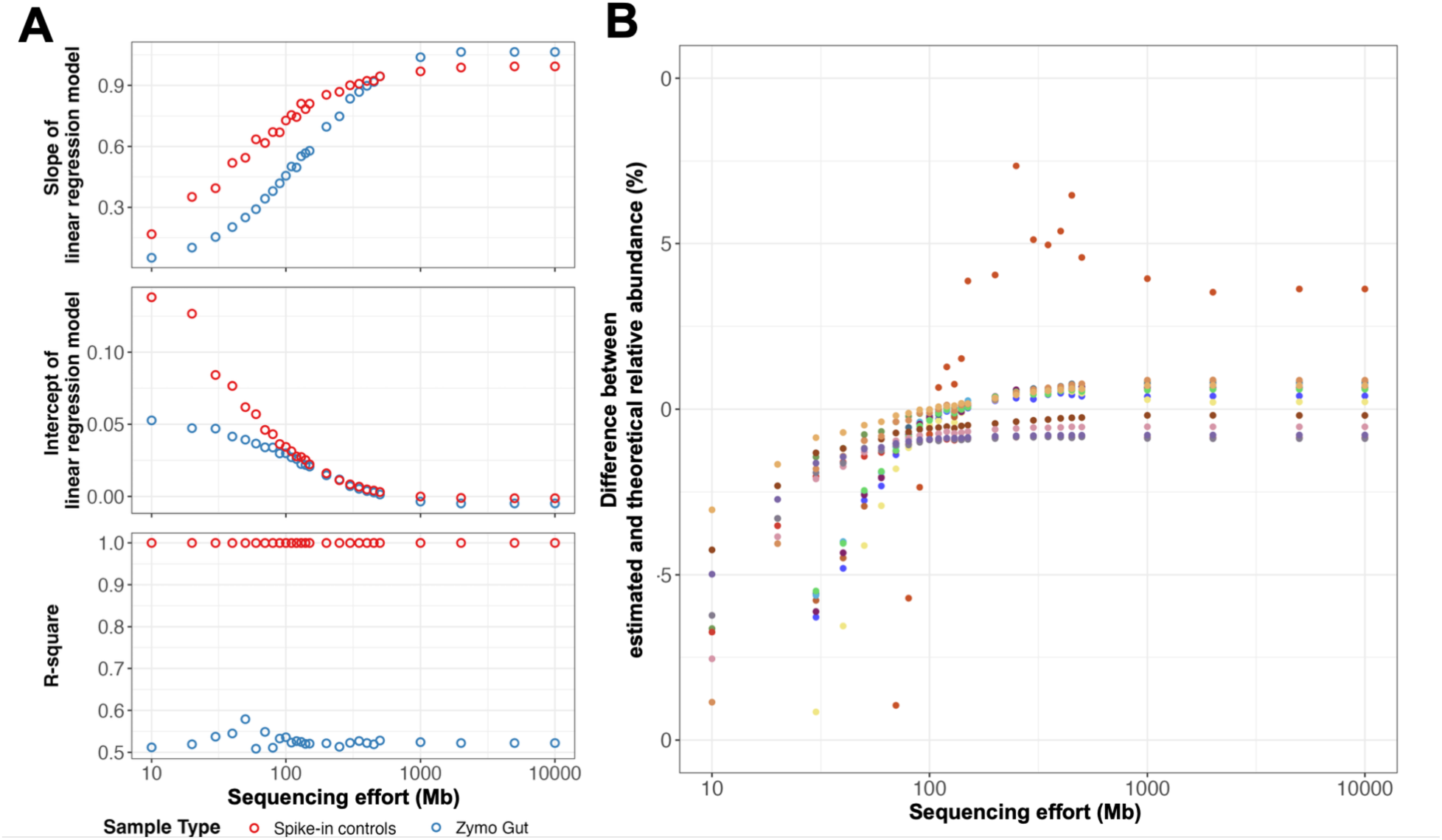
Comparison of coverage fraction and linear regression models between spike-in controls and mock samples (Zymo Gut). (A) Slope, intercept, and R^2^ values (correlation coefficient) of linear regression models for spike-in controls and ZG sample at different sequencing effort. Red and blue points represent data generated from spike-in controls and Zymo Gut, respectively. (B) Deviation of estimated relative abundance from theoretical values for each genome at different sequencing effort. The colors of the points represent different genomes, and the outlier corresponds to *Methanobrevibacter smithii*.

Here, it is important to note that the quantitation of genome copy numbers was based on the mapped bases of each genome (**Text S4, Eq. S5**) rather than the actual genome sizes (**Text S4, Eq. S6**)^30,54^. Actual genome sizes are often unavailable for microbial taxa due to incomplete genome assembly resulting from strain heterogeneity, repetitive elements, and low coverage of rare taxa^55–58^. BSINC strategy incorporates covered base-based calculation, thereby enabling quantitation of taxa with draft genomes, including eukaryotes^59^. This is particularly significant for environmental samples, where complete reference genomes are difficult to retrieve for the rare taxa present in complex communities^60^. Moreover, the BSINC strategy prevents cross-alignment between genomic spike-in control and sample reads by pre-sorting based on barcodes. While a set of synthetic DNA fragments (i.e., “sequins”) has been developed to represent a range of features and complexity, their distributions across attributes such as GC content and fragment length differ from those observed in natural microbial samples^32,61^. In contrast, the BSINC spike-in controls are derived from authentic microbial taxa and retain naturally occurring genomic characteristics, providing a more representative basis for quantitation than synthetic fragments^31,61,62^. Further, BSINC enables flexible optimization as different spike-in controls can be readily substituted to improve detection and quantitation accuracy.

The accuracy of absolute quantitation using the BSINC strategy was evaluated by comparing the relative abundance from estimated genome copy numbers against theoretical values for each genome at different sequencing efforts (**Figure 3B**). The difference between estimated relative abundance and theoretical values for each of the genomes in ZG samples decreased with increasing sequencing effort. However, DNA extraction biases also contribute to quantitative deviations, as similar relative abundance deviations are also observed in Illumina metagenomic sequencing (**Figure S8, Table S5**) and previous studies, even for high abundance taxa (e.g., *Roseburia hominis*, which accounts for 12.43% of genome copies in the community)^47^. Previous studies employed cellular-based spike-in controls added before DNA extraction to calculate DNA recovery rates and correct extraction biases^27,30^. However, each organism exhibits distinct cellular morphology and lysis susceptibility^47,63,64^. As linear regression models are designed to correct systematic biases that are consistent across organisms, the organism-dependent uncertainties in extraction biases may weaken the correlation between observed and theoretical genome copy numbers, reducing the accuracy of absolute quantitation^66^.

### 3.3. Establishing the dynamic quantitative limits of rD+rQ workflow

We established a limit of detection (LOD) as 10% of coverage fraction (i.e., at least 10% of a target genome should be covered by at least one sequenced read) as suggested by previous research^12,41,66^. Further, we adopted a dynamic approach for establishing limit of quantitation (LOQ) based on the acceptable coefficient of variation (CV) of spike-in controls and samples across replicates^35,36,62^. Coverage fraction and CVs of genome copy number were calculated for each genome in the spike-in controls and ZG samples (**Figures 4A** and **4B, Table S6)**. We set a 10% coverage fraction as the LOD and a 10% CV as the LOQ and non-detectable or non-quantifiable genomes were excluded. Then, a linear regression model was generated using the remaining genomes of spike-in controls (**Figure 4C**) and subsequently applied to estimate the observed genome copy number of ZG samples (**Figure 4D**). The estimated genome copy number was highly correlated with the theoretical genome copy number (**Figure 4D**, y = 0.9608x + 0.2719, R^2^=0.9645), indicating the accuracy of the rD+rQ workflow after incorporating an LOD and LOQ. Furthermore, dynamic LOD and LOQ adjustment based on sample-specific diversity allows optimization of sequencing effort during data acquisition with Nanopore sequencing. Previous research established a fixed LOD as the lowest input concentration of spike-in controls that was detected across all triplicates with acceptable CV with increasing sequencing efforts^33^. However, variable CVs were observed among taxa with same input concentrations within spike-in controls (e.g., *Cryptococcus neoformans* and *Enterococcus faecalis* in spike-in controls) or between spike-in controls and samples (e.g., *Pseudomonas aeruginosa* in spike-in controls and *Salmonella enterica* in samples), indicating that precision of quantitation was not solely determined by the concentration but also by taxon-specific characteristics (e.g., genome size, GC characteristics, community composition). Therefore, this fixed LOD is not transferable across different genomes.

**Figure 4.**
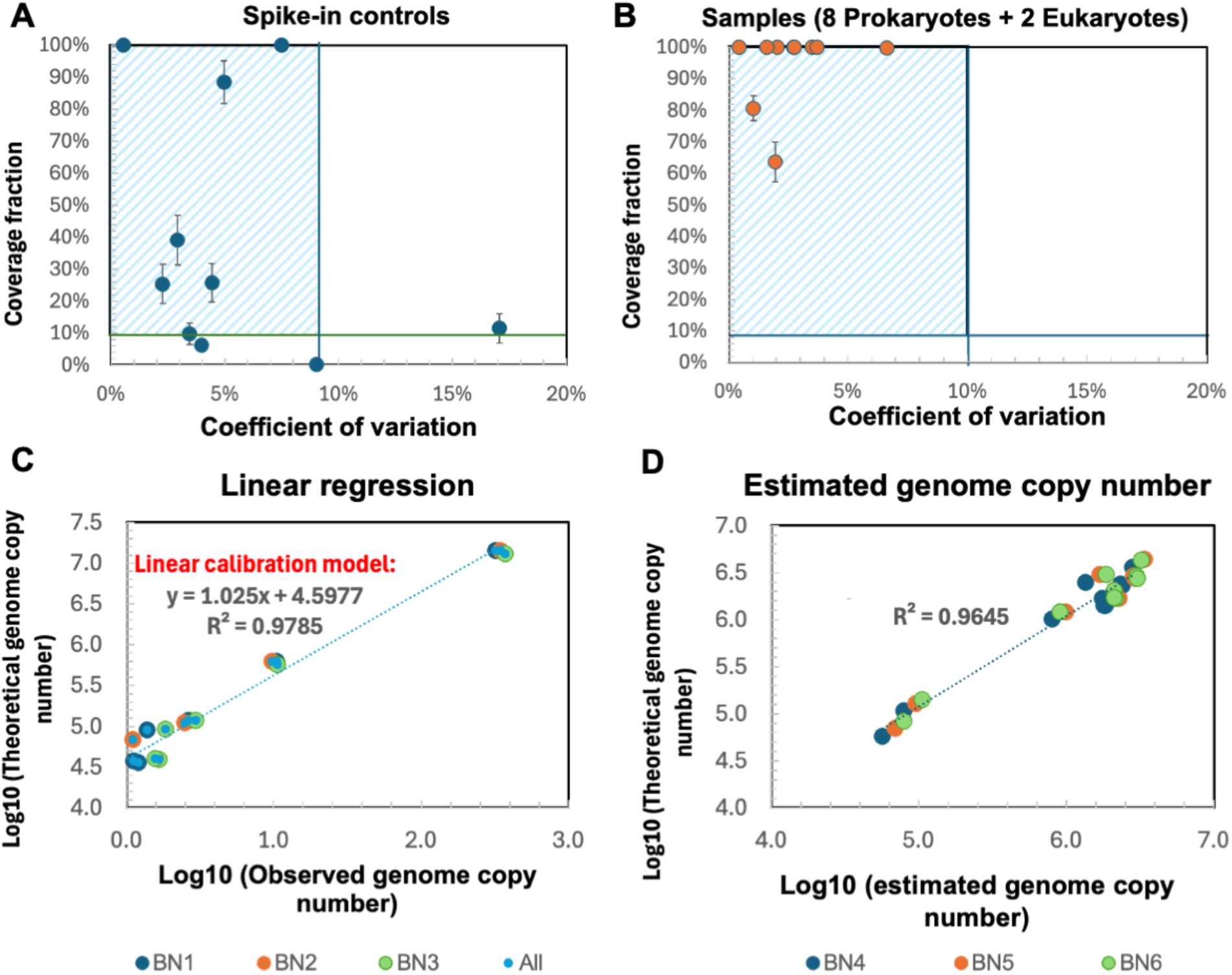
Coverage fraction and coefficient of variation (CV) of (A) Spike-in Controls and (B) ZG samples. Horizontal and vertical lines indicate the threshold of coverage fraction and the coefficient of variation, respectively. (C) Linear regression for log10-transformed theoretical genome copy number against observed genome copy numbers of spike-ins. (D) Correlation between log10-transformed theoretical genome copy number and estimated genome copy number of samples. The colors of the scatters indicated different barcodes (BN = Barcode number, All = Average of triplicates).

### 3.4. Utility of multiple taxa in spike-in controls compared to single spike-in control

To assess whether multiple-target spike-in controls, as discussed in section 3.3, provide improved quantitation accuracy compared to a single spike-in control, we estimated the genome copy numbers in ZG samples using a regression model derived from utilizing individual taxa in the spike-in control (i.e., single spike-in strategy) and compared to when the regression model was derived from all taxa in the spike-in control (i.e., rD+rQ approach). Estimated genome copy numbers using the rD+rQ approach exhibited significantly lower deviation from theoretical genome copy numbers as compared to the single spike-in in strategy for the ZG samples (**Figure 5, Table S7**). The microbial community in the spike-in control (Zymo Log) exhibits a log-distributed species abundance, with a significant disparity between dominant and rare species (Simpson index = 0.9, **Table S1).** In contrast, the ZG sample was relatively more even distributed (Simpson index = 0.11, **Table S3**), with comparable relative abundance across most taxa. Conventional strategies that provide single correction factor using single synthetic gene or genome spike-in control were less effective for calibration across all taxa in environmental samples^30,67^. As shown in **Figure 5**, substantial differences in abundance between single spike-in (e.g., *Staphylococcus aureus* (SA)) and sample taxa led to higher calibration errors (-96.91−7360.43%) than complex spike-ins (-23.60−95.26%). Some studies employed dual bacterial spike-in controls specifically to eliminate the extraction biases on different gram bacteria rather than to represent sample abundance diversity^67^. And the two bacteria were added at equal concentration and yielded single correction factors rather than regression models, limiting accurate estimation of taxa across varying abundance levels^30,67,68^.

**Figure 5.**
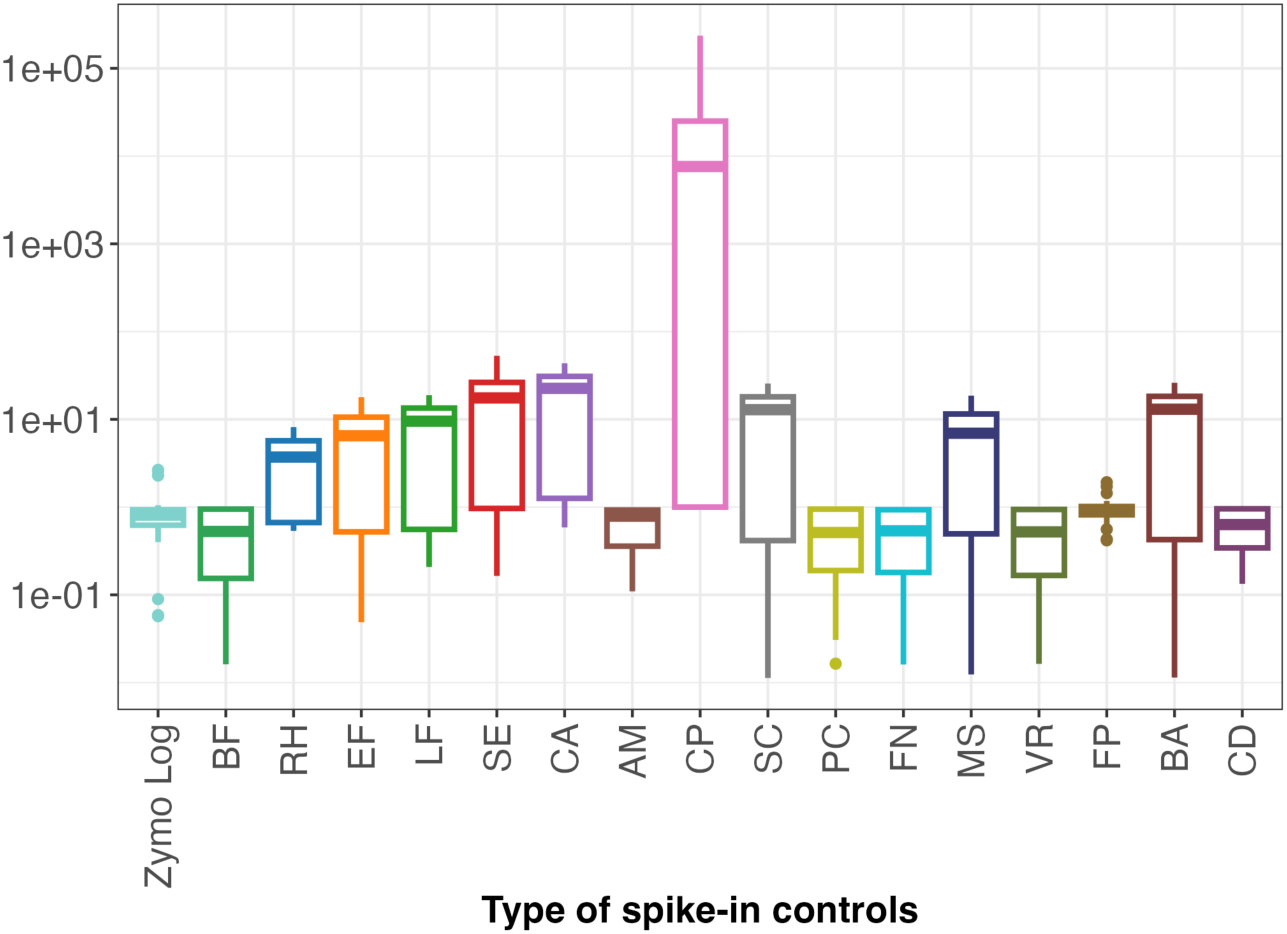
Comparison of absolute quantitation deviations of genomes in (A) Zymo Log and (B) Zymo Gut Communities resulting from different types of spike-in controls. On the x-axis, Zymo Log represents the complex spike-in controls used in the rD+rQ protocol, and the others are the abbreviations of single spike-in controls as listed in **Table S8**.

### 3.5. Benchmarking rD+rQ workflow in-field for environmental samples using digital PCR

Benchmarking to digital PCR (dPCR) was performed on ML and SE samples to determine the accuracy of quantitation using the portable rD+rQ workflow. To this end, we leveraged MAGs extracted from metagenomic data generated for ML and SE samples (**Text S3**) as the custom databases and nanopore sequencing data was mapped to these databases. Four genera, *Mycobacterium*, *Accumulibacter*, *Nitrospira*, and *Gordonia*, which were detected in both SE and ML samples were used as target genera for estimation of rD+rQ accuracy. There was a high agreement between the estimated genome copy numbers quantified using the rD+rQ workflow and dPCR for both ML and SE samples (**Figure 6**). Deviations in estimated abundance ranged from - 25.05% to -2.02% for *Mycobacterium*, *Accumulibacter* and *Gordonia* in ML samples and from - 0.04% to 23.99% for *Accumulibacter and Nitrospira* in SE samples. For the genus *Nitrospira* in ML samples and *Mycobacterium* and *Gordonia* in SE samples, rD+rQ workflow suggested that they were below the LOD (coverage fraction < 10%), explaining the high error. The dynamic rD+rQ workflow can address this limitation by flexible sequencing throughput based on requirements. When real-time results indicate taxa of interest below LOD or LOQ, total sequencing effort can be increased by continuing the sequencing, thereby improving analytical sensitivity for rare organisms^69,70^. Similarly, sequencing may be stopped and sequencing capacity (i.e., Flowcell pore viability) could be preserved once taxa of interest are determined to be above LOQ for accurate quantitation. Besides the low abundance, detection failures or underestimation could also be attributed to genomes that may be missing in the reference database. For instance, genus *Nitrospira* in ML samples (717.0 copies/µL DNA extract) and *Mycobacterium* in SE samples (1320.8 copies/µL DNA extract) were quantified by dPCR but remained below detection thresholds in rD+rQ analysis. This discrepancy could occur because the dPCR assay targets genus-level conserved regions that represent all species within the genus of interest, whereas rD+rQ requires species-level genomic matches for taxonomic assignment and aggregates individual species abundances to generate genus-level estimates^31^.

**Figure 6.**
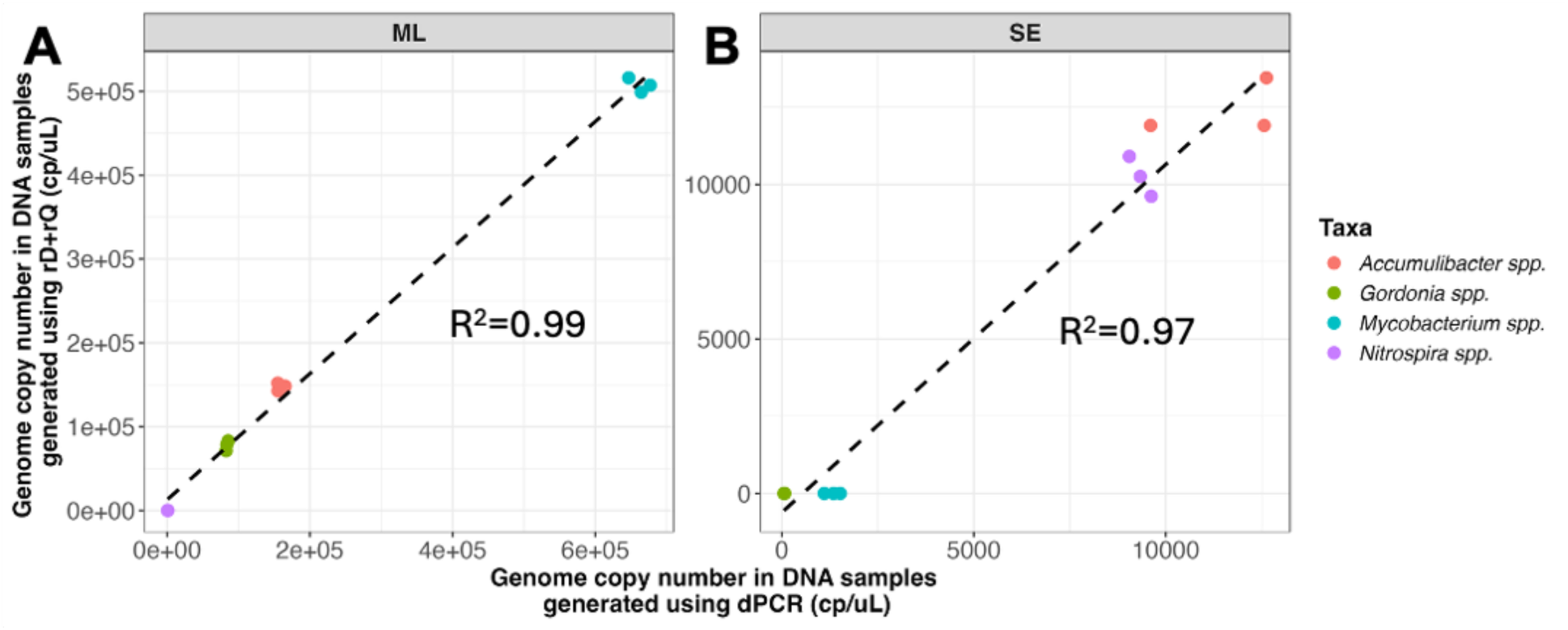
Comparison of Genome Copy Number Generated using digital PCR (dPCR) and Rapid Detection and Quantitation Workflow (rD+rQ) in mixed liquor sludge (ML) and secondary effluent (SE) samples, respectively.

## 4 Conclusion

This study developed a field-deployable Nanopore metagenomic sequencing-based workflow (rD+rQ workflow) and easy-to-use analysis pipeline based on EPI2ME for detection and absolute quantitation of microorganisms across various sample matrices in water sector. Immediate on-site processing preserves the original microbial community structure and nucleic acid integrity, effectively minimizes artifacts introduced by delayed processing, freezing–thawing cycles, or long-term storage. The BSINC strategy integrated within the rD+rQ workflow expands the application of natural genomic spike-ins for different types of samples, while enabling continuous optimization as new spike-in controls are developed. With dynamic LOD and LOQ, the rD+rQ workflow provides flexible solutions for real-time process control and water quality monitoring without predetermined thresholds, supporting adjustable and robust analysis for variable and unknown samples. However, this study has limitations in that DNA extraction biases cannot be eliminated. Further, quantitation of rare taxa may require extremely high sequencing effort which will increase associated costs. Moreover, robust taxonomic identification and quantitation depends on comprehensive reference databases. Future integration of selective Nanopore sequencing to enrich rare taxa may improve recovery of high-quality metagenome-assembled genomes from low-abundance species for the reference database.

## Data availability

The sequencing data for this project, including ZymoBIOMICS mock microbial community standards and environmental samples, is available in the NCBI under BioProject PRJNA1347130.

## Code availability

This simple-to-use detection and quantitation workflow incorporated with LOD and LOQ available on Github (https://github.com/Harper19/wf-metagenomics_test).

## Acknowledgments

The study was supported by The Water Research Foundation (Project 5100) and by the U.S. Department of Energy, Office of Energy Efficiency and Renewable Energy, under Award Number DE-EE0009270.

## References

1. Hatfield, R. G. et al. The Application of Nanopore Sequencing Technology to the Study of Dinoflagellates: A Proof of Concept Study for Rapid Sequence-Based Discrimination of Potentially Harmful Algae. Front. Microbiol. 11, (2020).

2. Werner, D. et al. MinION Nanopore Sequencing Accelerates Progress towards Ubiquitous Genetics in Water Research. Water 14, 2491 (2022).

3. Schurig, S. et al. Rapid Identification of Bacterial Composition in Wastewater by Combining Reverse Purification Nucleic Acid Extraction and Nanopore Sequencing. ACS EST Water 4, 1808–1818 (2024).

4. Latorre-Pérez, A., Pascual, J., Porcar, M. & Vilanova, C. A lab in the field: applications of real-time, in situ metagenomic sequencing. Biology Methods and Protocols 5, bpaa016 (2020).

5. Blackburn, J. et al. Use of synthetic DNA spike-in controls (sequins) for human genome sequencing. Nat Protoc 14, 2119–2151 (2019).

6. Handelsman, J. Metagenomics: Application of genomics to uncultured microorganisms. Microbiology and Molecular Biology Reviews 68, 669–685 (2004).

7. Eren, A. M. & Banfield, J. F. Modern microbiology: Embracing complexity through integration across scales. Cell 187, 5151–5170 (2024).

8. Ferreira, F. A., Helmersen, K., Visnovska, T., Jørgensen, S. B. & Aamot, H. V. Rapid nanopore-based DNA sequencing protocol of antibiotic-resistant bacteria for use in surveillance and outbreak investigation. Microbial Genomics 7, 000557 (2021).

9. Taş, N. et al. Metagenomic tools in microbial ecology research. Current Opinion in Biotechnology 67, 184–191 (2021).

10. Morgan, H. H., du Toit, M. & Setati, M. E. The Grapevine and Wine Microbiome: Insights from High-Throughput Amplicon Sequencing. Front. Microbiol. 8, (2017).

11. Yeluri Jonnala, B. R., McSweeney, P. L. H., Sheehan, J. J. & Cotter, P. D. Sequencing of the Cheese Microbiome and Its Relevance to Industry. Front. Microbiol. 9, (2018).

12. Lindner, B. G. et al. A user’s guide to the bioinformatic analysis of shotgun metagenomic sequence data for bacterial pathogen detection. International Journal of Food Microbiology 110488 (2023) doi:10.1016/j.ijfoodmicro.2023.110488.

13. Loose, M., Malla, S. & Stout, M. Real-time selective sequencing using nanopore technology. Nat Methods 13, 751–754 (2016).

14. Goenka, S. D. et al. Accelerated identification of disease-causing variants with ultra-rapid nanopore genome sequencing. Nat Biotechnol 40, 1035–1041 (2022).

15. Mitsuhashi, S. et al. A portable system for rapid bacterial composition analysis using a nanopore-based sequencer and laptop computer. Sci Rep 7, 5657 (2017).

16. Quick, J. et al. Real-time, portable genome sequencing for Ebola surveillance. Nature 530, 228–232 (2016).

17. Greninger, A. L. et al. Rapid metagenomic identification of viral pathogens in clinical samples by real-time nanopore sequencing analysis. Genome Medicine 7, 99 (2015).

18. Buytaers, F. E. et al. Towards Real-Time and Affordable Strain-Level Metagenomics-Based Foodborne Outbreak Investigations Using Oxford Nanopore Sequencing Technologies. Frontiers in Microbiology 12, (2021).

19. Branders, S., Grabherr, M. G. & Ahmad, R. Real-time Taxonomic Characterization of Long-read Mixed-species Sequencing Samples in Sorted Motif Distance Space: Voyager. 2024.04.13.589333 Preprint at 10.1101/2024.04.13.589333 (2024).

20. Knudsen, B. E., et al. Impact of Sample Type and DNA Isolation Procedure on Genomic Inference of Microbiome Composition. mSystems 1, 10.1128/msystems.00095-16 (2016).

21. Schurig, S. et al. Rapid Identification of Bacterial Composition in Wastewater by Combining Reverse Purification Nucleic Acid Extraction and Nanopore Sequencing. ACS EST Water 4, 1808–1818 (2024).

22. Maghini, D. G., Moss, E. L., Vance, S. E. & Bhatt, A. S. Improved high-molecular-weight DNA extraction, nanopore sequencing and metagenomic assembly from the human gut microbiome. Nat Protoc 16, 458–471 (2021).

23. Karstens, L. et al. Benchmarking DNA isolation kits used in analyses of the urinary microbiome. Sci Rep 11, 6186 (2021).

24. Mason, M. G. & Botella, J. R. Rapid (30-second), equipment-free purification of nucleic acids using easy-to-make dipsticks. Nat Protoc 15, 3663–3677 (2020).

25. Martello, A., Lambert, B., Johnston, C., Cutler, J. & Stumpf, C. F. Comparison of the novel dipstick DNA extraction technique with two established techniques for use in biological barcoding. Mol Biol Rep 46, 6625–6628 (2019).

26. Irwin, P. et al. The near-quantitative sampling of genomic DNA from various food-borne Eubacteria. BMC Microbiol 14, 326 (2014).

27. Barlow, J. T., Bogatyrev, S. R. & Ismagilov, R. F. A quantitative sequencing framework for absolute abundance measurements of mucosal and lumenal microbial communities. Nat Commun 11, 2590 (2020).

28. Reis, A. L. M. et al. A universal and independent synthetic DNA ladder for the quantitative measurement of genomic features. Nat Commun 11, 3609 (2020).

29. Deveson, I. W. et al. Representing genetic variation with synthetic DNA standards. Nat Methods 13, 784–791 (2016).

30. Yang, Y. et al. Rapid absolute quantification of pathogens and ARGs by nanopore sequencing. Science of The Total Environment 809, 152190 (2022).

31. Davis, B. C., Vikesland, P. J. & Pruden, A. Evaluating Quantitative Metagenomics for Environmental Monitoring of Antibiotic Resistance and Establishing Detection Limits. Environ. Sci. Technol. 59, 6192–6202 (2025).

32. Langenfeld, K. et al. A quantitative metagenomic approach to determine population concentrations with examination of quantitative limitations. 2022.07.08.499345 Preprint at 10.1101/2022.07.08.499345 (2022).

33. Davis, B. C., Vikesland, P. J. & Pruden, A. Evaluating Quantitative Metagenomics for Environmental Monitoring of Antibiotic Resistance and Establishing Detection Limits. Environ. Sci. Technol. (2025) doi:10.1021/acs.est.4c08284.

34. Bradley, I. M., Pinto, A. J. & Guest, J. S. Design and Evaluation of Illumina MiSeq-Compatible, 18S rRNA Gene-Specific Primers for Improved Characterization of Mixed Phototrophic Communities. Applied and Environmental Microbiology 82, 5878–5891 (2016).

35. Alam, M. M. et al. Community Structure and Function During Periods of High Performance and System Upset in a Full-Scale Mixed Microalgal Wastewater Resource Recovery Facility. 2024.01.23.576871 Preprint at 10.1101/2024.01.23.576871 (2024).

36. Callahan, B. J. et al. DADA2: High-resolution sample inference from Illumina amplicon data. Nat Methods 13, 581–583 (2016).

37. Quast, C. et al. The SILVA ribosomal RNA gene database project: improved data processing and web-based tools. Nucleic Acids Res 41, D590–D596 (2013).

38. Yilmaz, P. et al. The SILVA and “All-species Living Tree Project (LTP)” taxonomic frameworks. Nucleic Acids Res 42, D643–D648 (2014).

39. Davis, N. M., Proctor, D. M., Holmes, S. P., Relman, D. A. & Callahan, B. J. Simple statistical identification and removal of contaminant sequences in marker-gene and metagenomics data. Microbiome 6, 226 (2018).

40. Wickham, H. Data Analysis. in *ggplot2: Elegant Graphics for Data Analysis* (ed. Wickham, H.) 189–201 (Springer International Publishing, Cham, 2016). doi:10.1007/978-3-319-24277-4_9.

41. Castro, J. C. et al. imGLAD: accurate detection and quantification of target organisms in metagenomes. PeerJ 6, e5882 (2018).

42. Aroney, S. T. N. et al. CoverM: read alignment statistics for metagenomics. Bioinformatics 41, btaf147 (2025).

43. Radomski, N., Kreitmann, L., McIntosh, F. & Behr, M. A. The Critical Role of DNA Extraction for Detection of Mycobacteria in Tissues. PLOS ONE 8, e78749 (2013).

44. He, S., Gall, D. L. & McMahon, K. D. “Candidatus Accumulibacter” Population Structure in Enhanced Biological Phosphorus Removal Sludges as Revealed by Polyphosphate Kinase Genes. Appl Environ Microbiol 73, 5865–5874 (2007).

45. Vilardi, K. J. et al. Nitrogen source influences the interactions of comammox bacteria with aerobic nitrifiers. Microbiology Spectrum 12, e03181–23 (2024).

46. Xu, S., Sun, M., Zhang, C., Surampalli, R. & Hu, Z. Filamentous sludge bulking control by nano zero-valent iron in activated sludge treatment systems. Environ. Sci.: Processes Impacts 16, 2721–2728 (2014).

47. Pu, Y. et al. Impact of DNA Extraction Methods on Gut Microbiome Profiles: A Comparative Metagenomic Study. Phenomics 5, 76–90 (2025).

48. Shi, Z. et al. The Effects of DNA Extraction Kits and Primers on Prokaryotic and Eukaryotic Microbial Community in Freshwater Sediments. Microorganisms 10, 1213 (2022).

49. Bramucci, A. R. et al. Microvolume DNA extraction methods for microscale amplicon and metagenomic studies. ISME COMMUN. 1, 1–5 (2021).

50. Albertsen, M., Karst, S. M., Ziegler, A. S., Kirkegaard, R. H. & Nielsen, P. H. Back to Basics – The Influence of DNA Extraction and Primer Choice on Phylogenetic Analysis of Activated Sludge Communities. PLOS ONE 10, e0132783 (2015).

51. Guo, F. & Zhang, T. Biases during DNA extraction of activated sludge samples revealed by high throughput sequencing. Appl Microbiol Biotechnol 97, 4607–4616 (2013).

52. Acharya, K. et al. Metagenomic water quality monitoring with a portable laboratory. Water Research 184, 116112 (2020).

53. Zhang, L. et al. Comparison Analysis of Different DNA Extraction Methods on Suitability for Long-Read Metagenomic Nanopore Sequencing. Front. Cell. Infect. Microbiol. 12, (2022).

54. Langenfeld, K. et al. Development of a quantitative metagenomic approach to establish quantitative limits and its application to viruses. 2022.07.08.499345 Preprint at 10.1101/2022.07.08.499345 (2024).

55. Albertsen, M. et al. Genome sequences of rare, uncultured bacteria obtained by differential coverage binning of multiple metagenomes. Nat Biotechnol 31, 533–538 (2013).

56. Zhang, Z. et al. Exploring high-quality microbial genomes by assembling short-reads with long-range connectivity. Nat Commun 15, 4631 (2024).

57. Bouras, G. et al. How low can you go? Short-read polishing of Oxford Nanopore bacterial genome assemblies. Microbial Genomics 10, 001254 (2024).

58. Zeng, S. et al. A compendium of 32,277 metagenome-assembled genomes and over 80 million genes from the early-life human gut microbiome. Nat Commun 13, 5139 (2022).

59. Ayling, M., Clark, M. D. & Leggett, R. M. New approaches for metagenome assembly with short reads. Brief Bioinform 21, 584–594 (2020).

60. Loeffler, C. et al. Improving the usability and comprehensiveness of microbial databases. BMC Biology 18, 37 (2020).

61. Blackburn, J. et al. Use of synthetic DNA spike-in controls (sequins) for human genome sequencing. Nat Protoc 14, 2119–2151 (2019).

62. Li, B., Li, X. & Yan, T. A Quantitative Metagenomic Sequencing Approach for High-Throughput Gene Quantification and Demonstration with Antibiotic Resistance Genes. Applied and Environmental Microbiology 87, e00871–21 (2021).

63. Feinstein, L. M., Sul, W. J. & Blackwood, C. B. Assessment of Bias Associated with Incomplete Extraction of Microbial DNA from Soil. Applied and Environmental Microbiology 75, 5428–5433 (2009).

64. Crossette, E. et al. Metagenomic Quantification of Genes with Internal Standards. mBio 12, 10.1128/mbio.03173-20 (2021).

65. Hardwick, S. A. et al. Synthetic microbe communities provide internal reference standards for metagenome sequencing and analysis. Nat Commun 9, 3096 (2018).

66. Lindner, B. G. et al. Toward shotgun metagenomic approaches for microbial source tracking sewage spills based on laboratory mesocosms. Water Research 210, 117993 (2022).

67. Yang, Y. et al. QMRA of beach water by Nanopore sequencing-based viability-metagenomics absolute quantification. Water Research 235, 119858 (2023).

68. Camacho-Sanchez, M. A new spike-in-based method for quantitative metabarcoding of soil fungi and bacteria. Int Microbiol 27, 719–730 (2024).

69. Sims, D., Sudbery, I., Ilott, N. E., Heger, A. & Ponting, C. P. Sequencing depth and coverage: key considerations in genomic analyses. Nat Rev Genet 15, 121–132 (2014).

70. Gao, W. et al. Low-depth Raw Reads of Nanopore Sequencing Enables Rapid and Accurate Bacterial Identification. Appl Biochem Microbiol 59, 968–974 (2023).

