## Supplemental information for "Quantitative metagenomics using a portable protocol"

**Affiliations of authors:**

^a^ School of Civil and Environmental Engineering, Georgia Institute of Technology, Atlanta, GA 30332, USA

^b^ Great Lakes Water Authority, Detroit, MI 48209, USA

^c^ Hampton Roads Sanitation District, Virginia Beach, VA 23455, USA

^d^ Hazen and Sawyer, Fairfax, VA 22030, USA

^e^ School of Earth and Atmospheric Sciences, Georgia Institute of Technology, Atlanta, GA 30332, USA

**Total pages: 23**

**Number of texts: 4**

**Number of figures: 8**

**Number of tables: 8**

**Contents**

**Text S1.** Optimization of SuperFastPrep-2 power settings.

The speed of lysis controlled by the power level of the rotary tool in the SuperFastPrep-2 was optimized. As shown in **Figure S1**, the fragment size decreased with higher power levels. The DNA yield increased at higher power levels until power level 20 after which it decreased. This suggests that DNA was likely over-sheared at power level 25, resulting in the loss of short DNA fragments during the subsequent purification steps. Considering the compromise between DNA yield and shearing, both power levels 15 and 20, which exhibited performance comparable to the conventional benchtop homogenizer FastPrep-24, were employed for further evaluation.

**Test S2.** Definition of Key Terms Related to Genomic Analysis

Covered bases: The number of bases in a reference genome that are covered by at least one sequence read.

Mapped bases: The number of bases from the sequencing reads that are aligned (mapped) to a reference genome.

Sequencing depth: The average number of sequenced reads across all base pair positions of a genome.

Coverage fraction: The fraction of base pair positions in a genome covered by at least one sequence read.

Sequencing effort: The amount of data produced by the sequencing run.

**Text S3.** Development of custom reference genomic database.

Samples were collected from relevant water utilities and subjected to DNA extraction using DNeasy PowerWater following the manufacturer’s protocol, followed by metagenomic sequencing at Georgia Institute of Technology’s Molecular Evolution Core on the Illumina HiSeq sequencing platform. A total of 20 GB of 150-bp paired-end data was retrieved from each sample and adapters and low-quality sequences were trimmed and filtered using fastp v0.23.4^1^. Vector contamination was detected and removed based on UniVec Core 10.0 database using SAMtools^2–5^. Subsequently, a single sample assembly was performed on the data of each sample using MetaSPAdes^6^. Single sample assemblies were binned with MetaBAT2 (using contigs larger than 2000 bp), VAMB (using contigs larger than 2000 bp), and SemiBin2^7–9^. The results were combined using DASTools to generate an optimized and non-redundant set of genome bins^10^. The quality assessment for generated genome bins is shown in Figure 4-1. Medium (50 < completeness ≤ 95, 5 < contamination ≤ 10) and high (completeness ≥ 95, contamination ≤ 5) quality genomes were retained in the system-specific reference databases.

**Text S4.** Equations.

${BEF}_{i}= \frac{Total DNA input of {barcode}_{i} \left( ng \right)\times Total sequencing effort of {barcode}_{1} (bp)}{Total sequencing effort of {barcode}_{i} \left( bp \right) \times Total DNA input of {barcode}_{1} \left( ng \right)}$	Eq. S1

$BEF normalized mapped bases of {genome}_{j} in {barcode}_{i}=Observed mapped bases of {genome}_{j} in{barcode}_{i} \times{BEF}_{i}$	Eq. S2

$Observed copy number of {genome}_{j} in {barcode}_{i} = \frac{BEF normalized mapped bases of {genome}_{j} in {barcode}_{i}}{Covered bases of {genome}_{j} in {barcode}_{i}}$	Eq. S3

$Deviation of genome copy number = \frac{Calibrated genome copy number of genome-Theoretical genome copy number}{Theoretical genome copy number}$	Eq. S4

$Genome copy number= \frac{Mapped bases of the taxon (bp)}{Covered bases (bp)}$	Eq. S5

$Sequencing effots per taxon (mean depth)= \frac{Mapped bases of the taxon (bp)}{Genome size of the taxon (bp)}$	Eq. S6

Where i is the ID of the barcode ligated to the sample and barcode_1_ is one of the spike-in controls. All the other barcodes were compared to the barcode_1_. Genome_j_ represents different unique genome.

### Figures

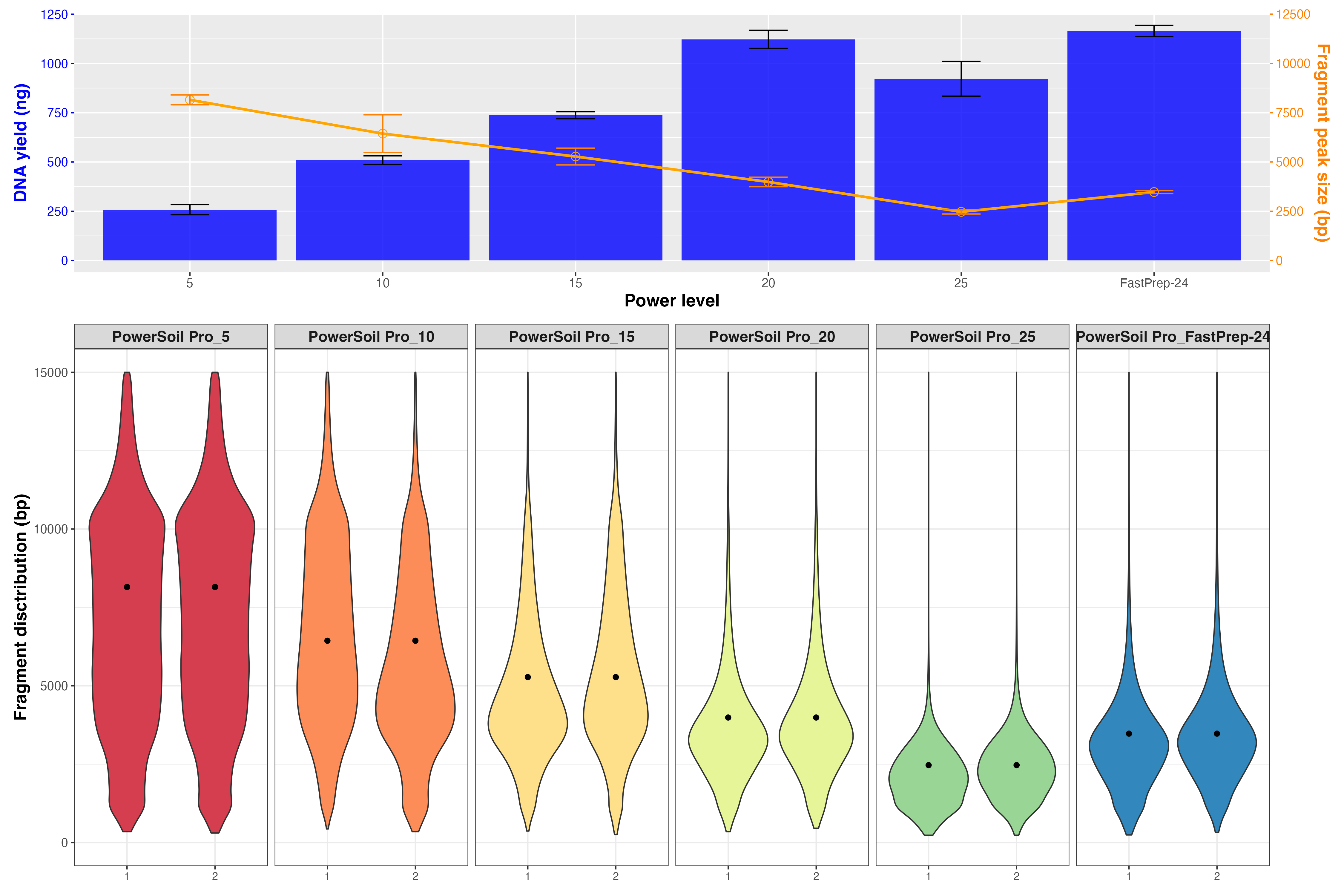

**Figure S1.** Impact of different lysis power levels of handheld bead beater. (A) DNA yield (blue bars) and fragment peak size (orange line), and (B) fragment distribution (violin plot) and peak size (filled circle) for different power levels (5-25) of the handheld bead beater utilized for bead beating. 1 and 2 on the x-axis represent replicate extraction. The FastPrep-24, a widely utilized bench-scale device for bead beating in the laboratory, is used as a reference for comparison.

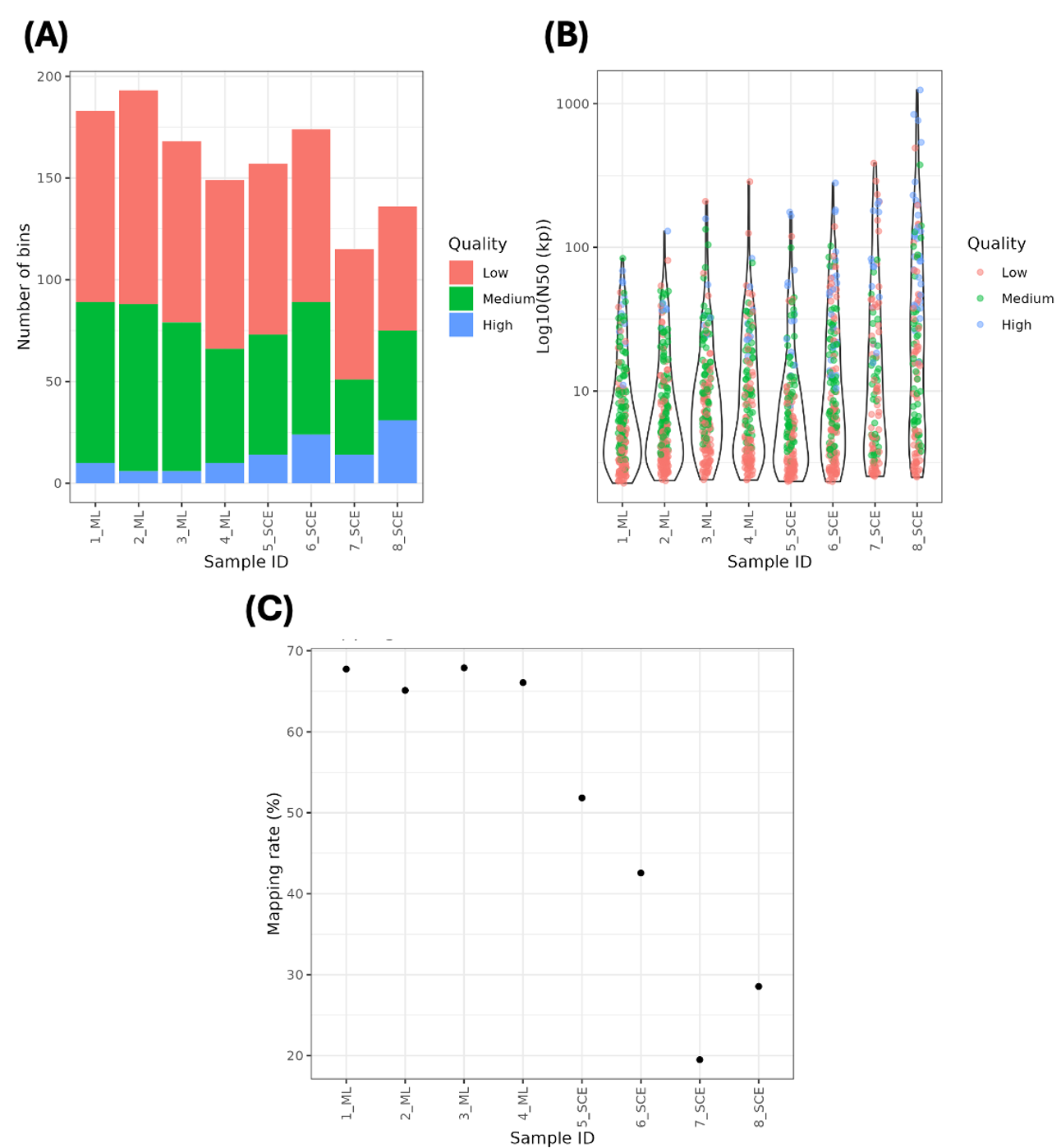

**Figure S2.** Summary of databases generated from different water facilities. (A) Number and (B) contiguity (i.e., N50) of bins of different levels of quality in terms of completeness and contamination. (C) Proportion of metagenomic reads of samples that were mapped against the draft bins. High, medium, and low indicate the different quality levels. Low: completeness < 50%, contamination > 10%; Medium: 50% < completeness ≤ 95%, 5% < contamination ≤ 10%; High: completeness ≥ 95%, contamination ≤ 5%.

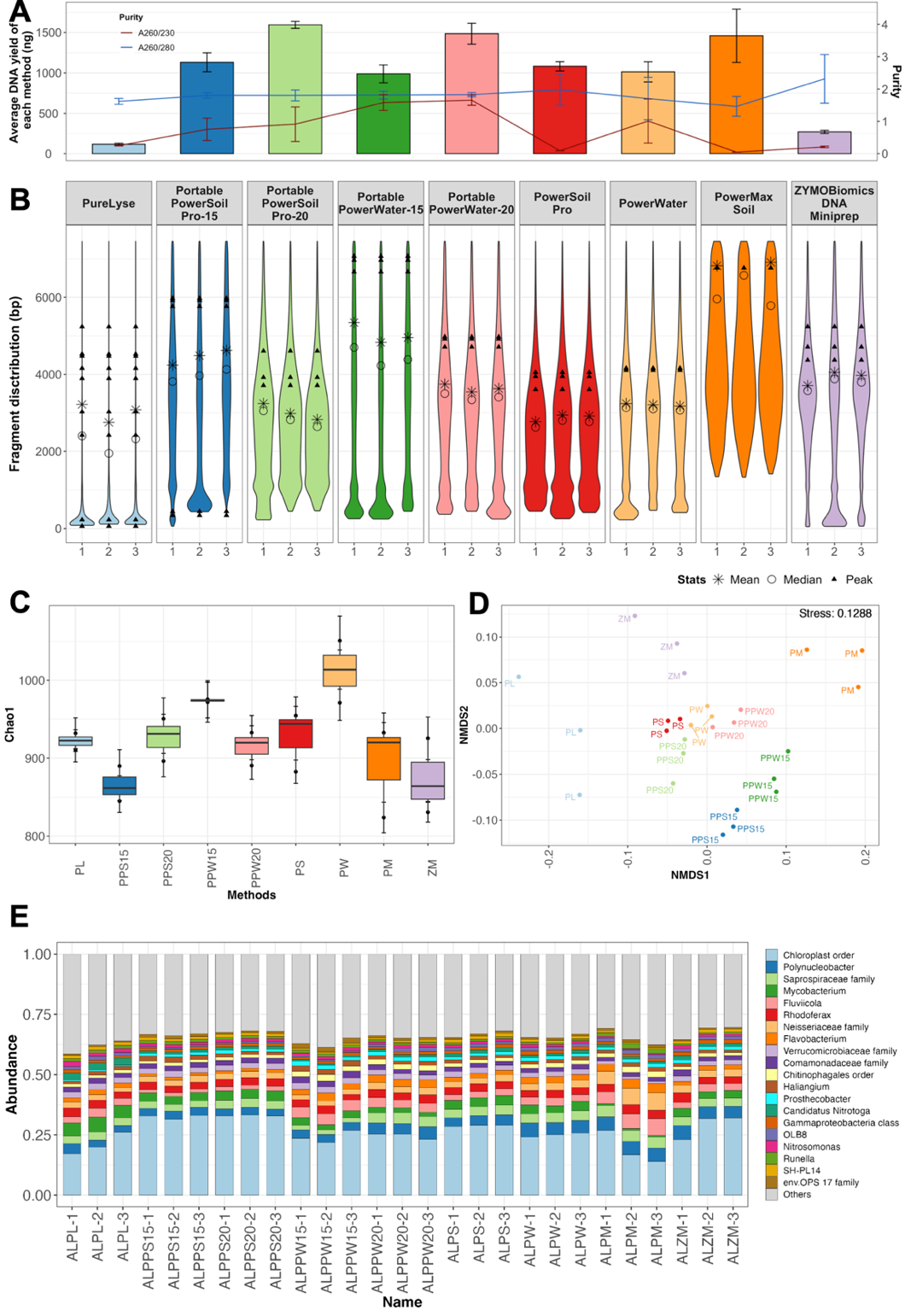

**Figure S3.** Evaluation of DNA extraction methods on the microbial community in mix tank effluent samples (AL). (A) Purity and quantity, and (B) DNA fragment size distribution for DNA extracted using different protocols. The line, bar, and bubble plots indicate the average purity, DNA yield, and fragment distribution, respectively. The star, circle, and filled triangle up represent the mean, median, and peak of DNA fragments by molarity, respectively. (C) Alpha diversity (Chao1 index), (D) Beta diversity, and (E) composition of the microbial community for prokaryotes. The Beta diversity is measured using NMDS2 plot based on Bray-Curtis distance.

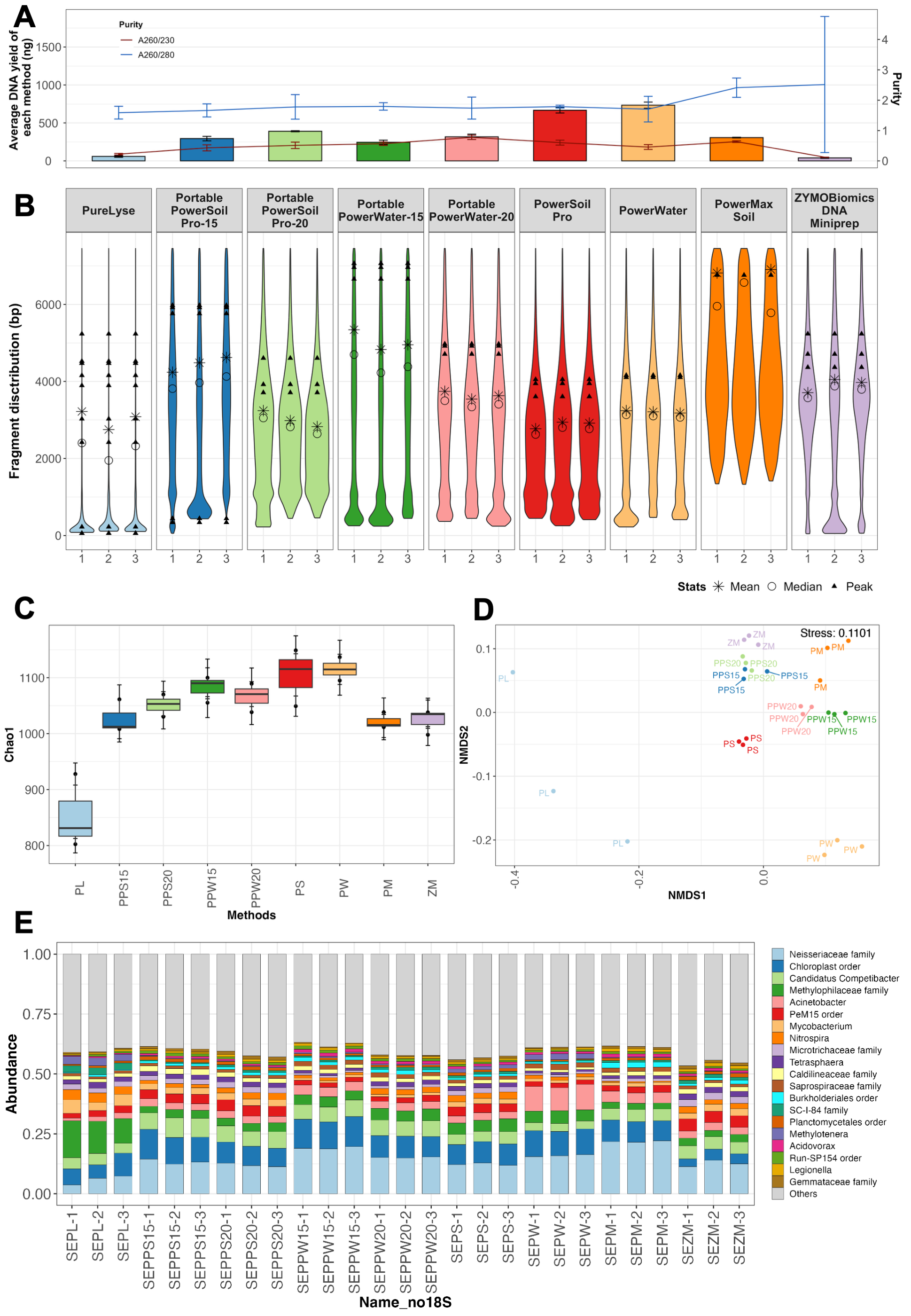

**Figure S4.** Evaluation of DNA extraction methods on the microbial community in secondary effluent samples (SE). (A) Purity and quantity, and (B) fragment distribution of DNA extracted uinsg different protocols. The line, bar, and bubble plots indicate the average purity, DNA yield, and fragment distribution, respectively. The star, circle, and filled triangle up represent the mean, median, and peak of DNA fragments by molarity, respectively. (C) Alpha diversity (Chao1 index), (D) Beta diversity, and (E) composition of the microbial community for prokaryotes. The Beta diversity is measured using NMDS2 plot based on Bray-Curtis distance.

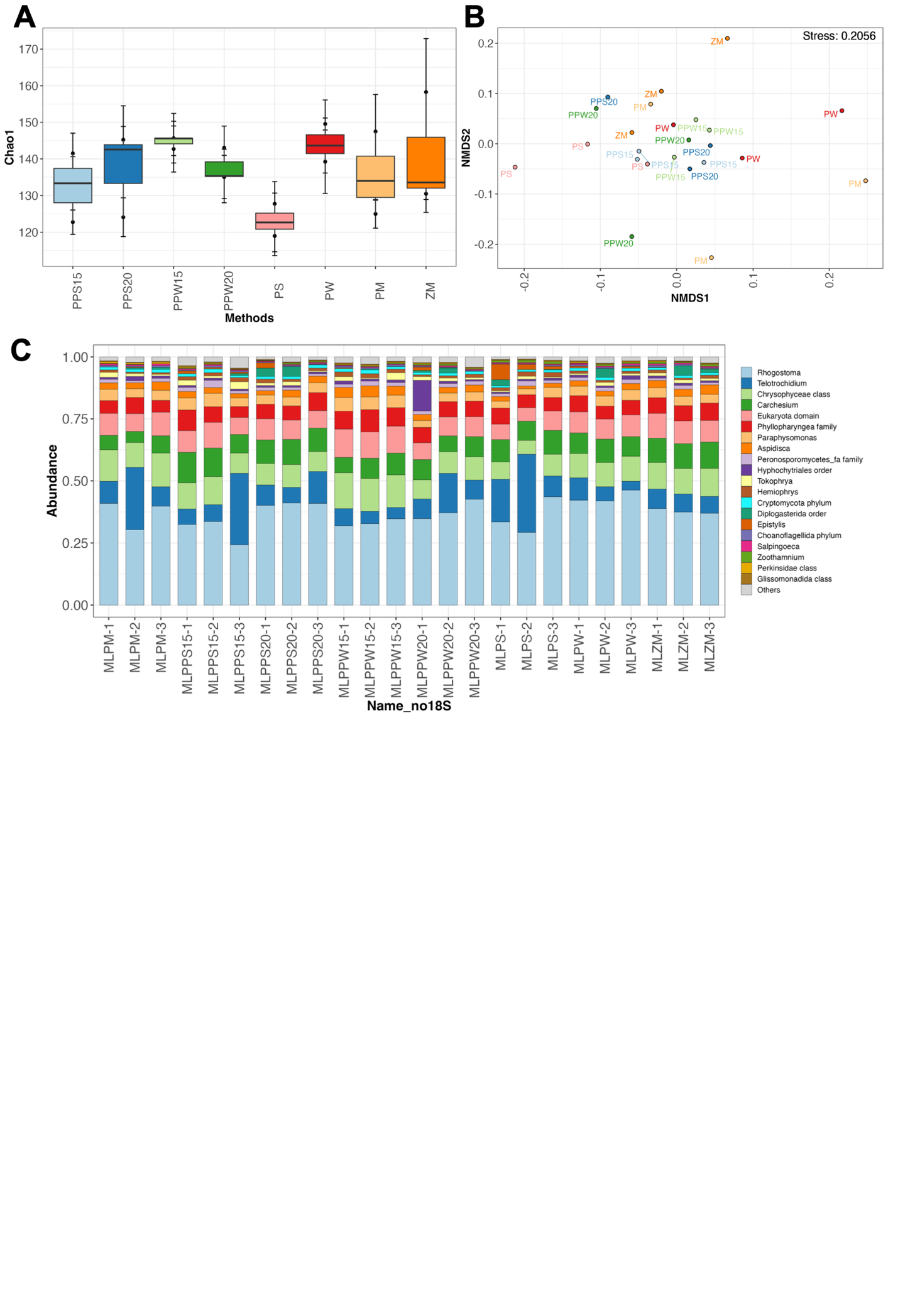

**Figure S5.** Evaluation of DNA extraction methods on the eukaryotic community in mixed liquor sludge samples (ML). (A) Alpha diversity (Chao1 index), (B) Beta diversity, and (C) composition of the microbial community for eukaryotes. The Beta diversity is measured using NMDS2 plot based on Bray-Curtis distance.

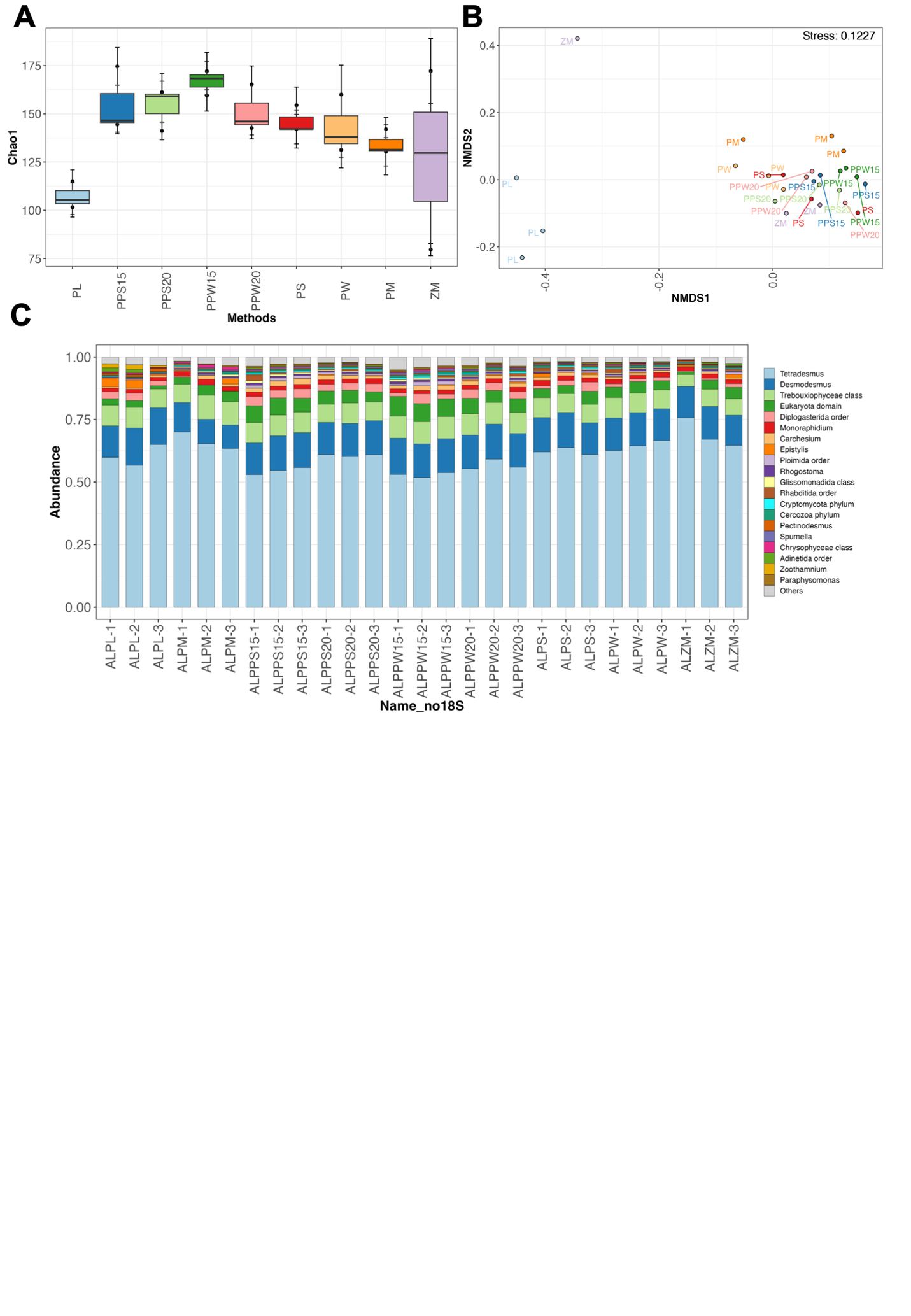
 **Figure S6.** Evaluation of DNA extraction methods on the eukaryotic community in mix tank effluent samples (AL). (A) Alpha diversity (Chao1 index), (B) Beta diversity, and (C) composition of the microbial community for eukaryotes. The Beta diversity is measured using NMDS2 plot based on Bray-Curtis distance.

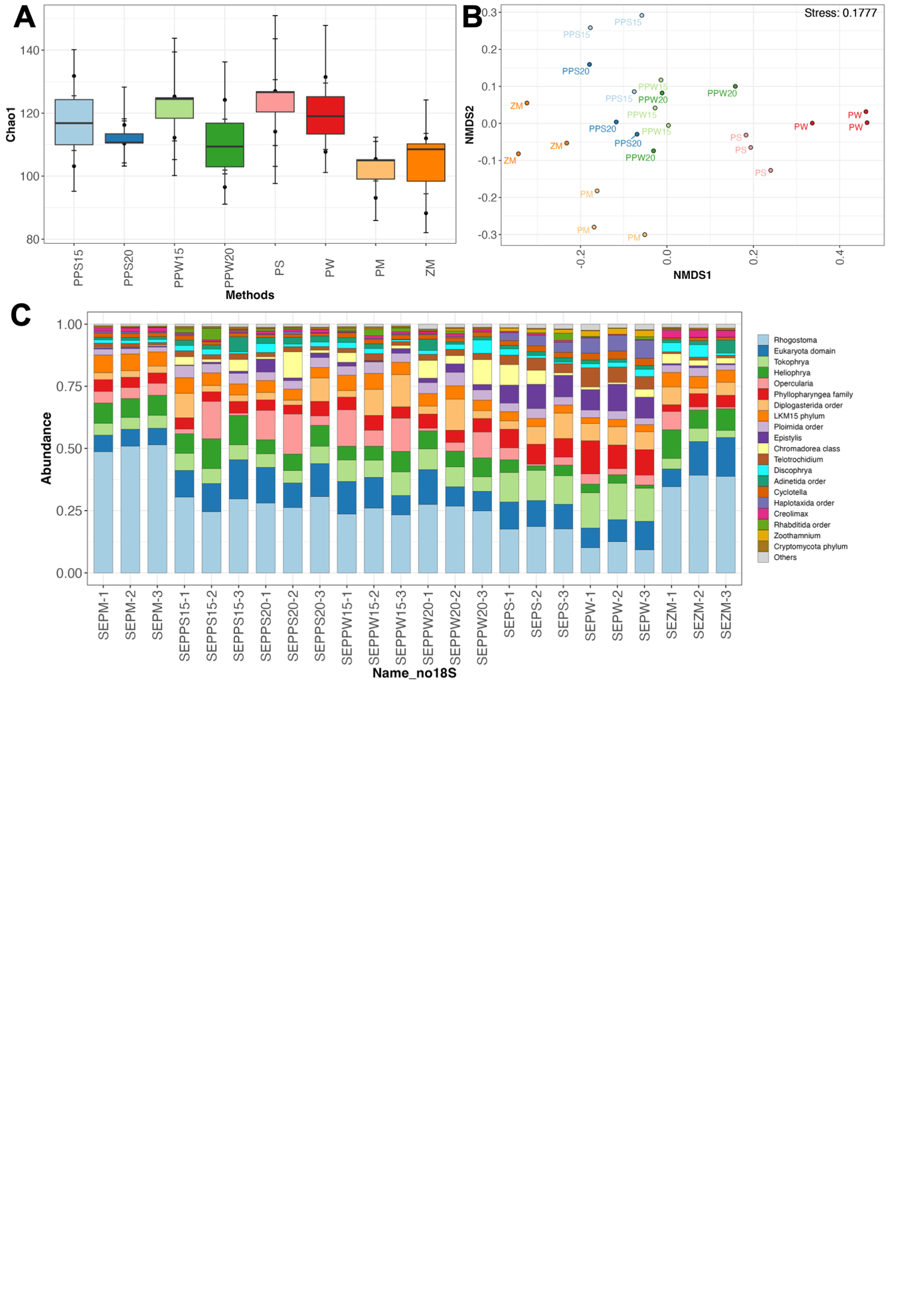

**Figure S7.** Evaluation of DNA extraction methods on the eukaryotic community in secondary effluent samples (SE). (A) Alpha diversity (Chao1 index), (B) Beta diversity, and (C) composition of the microbial community for *eukaryotes*. The Beta diversity is measured using NMDS2 plot based on Bray-Curtis distance.

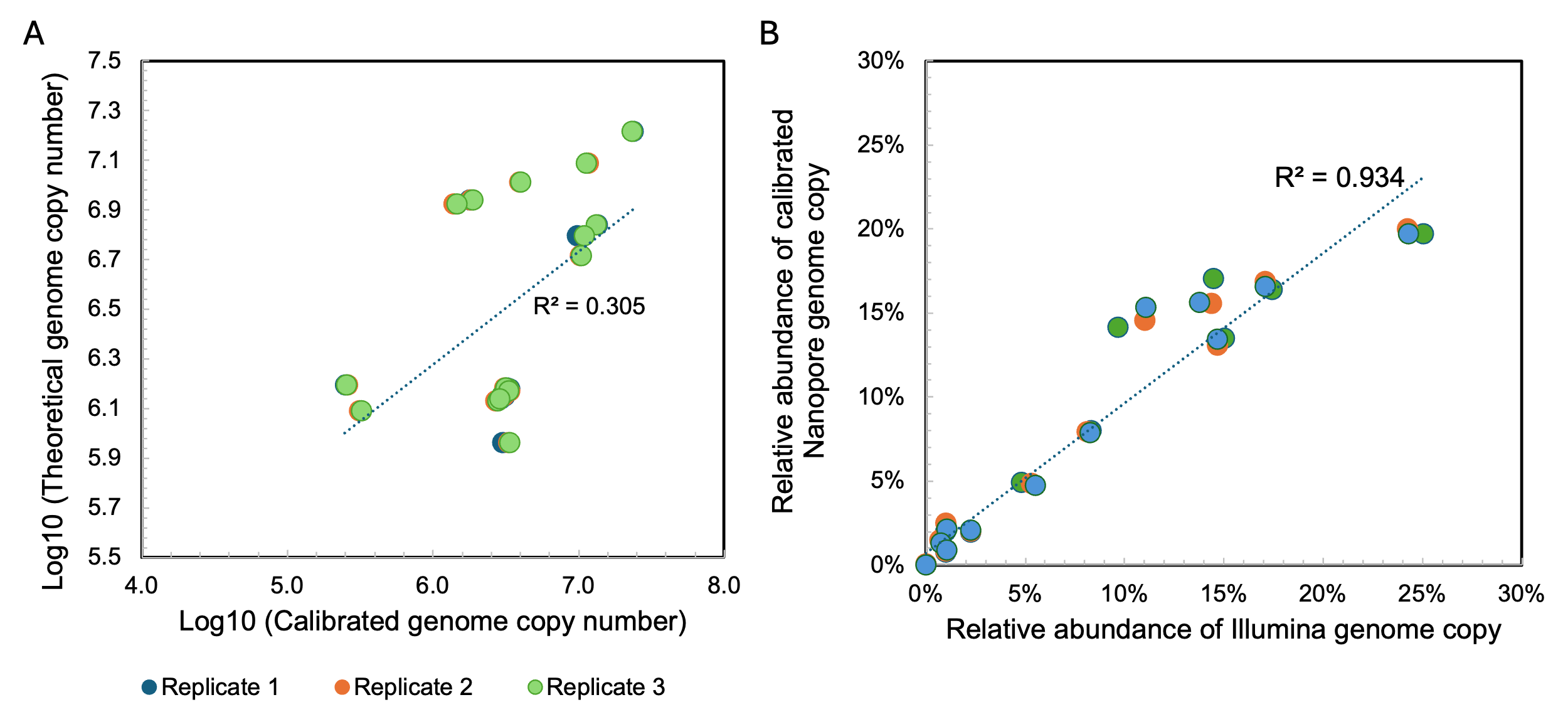

**Figure S8.** Results of Absolute Quantitation in ZYMO Gut Community using rD+rQ Protocol. (A) Correlation between log10-transformed theoretical genome copy number against observed genome copy numbers of samples. (B) Correlation between the relative abundance of genome copy number of Nanopore against Illumina data. The colors of the scatters indicated triplicates of each genome.

**Tables.**

**Table S1.** Composition of ZymoBIOMICS Community DNA Standard II (Log Distribution).

| Abbreviation | Taxa | Theoretical Composition (%) | | | | | |
| --- | --- | --- | --- | --- | --- | --- | --- |
|  |  | Genomic DNA | 16S only | 16S & 18S | Genome Copy | Cell Number | Gram Stain |
| SA | *Staphylococcus aureus* | 0.000089 | 0.0001 | 0.0001 | 0.0001 | 0.0001 | + |
| CN | *Cryptococcus neoformans* | 0.00089 | NA | 0.0014 | 0.00015 | 0.00007 | Yeast |
| EF | *Enterococcus faecalis* | 0.00089 | 0.00067 | 0.00064 | 0.001 | 0.001 | + |
| LF | *Lactobacillus fermentum* | 0.0089 | 0.012 | 0.012 | 0.015 | 0.015 | + |
| EC | *Escherichia coli* | 0.089 | 0.069 | 0.066 | 0.058 | 0.058 | - |
| SE | *Salmonella enterica* | 0.089 | 0.07 | 0.067 | 0.059 | 0.059 | - |
| BS | *Bacillus subtilis* | 0.89 | 1.2 | 1.1 | 0.7 | 0.7 | + |
| SC | *Saccharomyces cerevisiae* | 0.89 | NA | 4.1 | 0.23 | 0.12 | Yeast |
| PA | *Pseudomonas aeruginosa* | 8.9 | 2.8 | 2.7 | 4.2 | 4.2 | - |
| LM | *Listeria monocytogenes* | 89.1 | 95.9 | 91.9 | 94.8 | 94.9 | + |

**Table S2**. Composition of ZymoBIOMICS Microbial Community Standard (Even Distribution).

| Abbreviation | Species | Theoretical Composition (%) | | | | |
| --- | --- | --- | --- | --- | --- | --- |
|  |  | Genomic DNA | 16S only | 16S & 18S | Genome Copy | Cell Number |
| SA | *Pseudomonas aeruginosa* | 12 | 4.2 | 3.6 | 6.1 | 6.1 |
| CN | *Escherichia coli* | 12 | 10.1 | 8.9 | 8.5 | 8.5 |
| EF | *Salmonella enterica* | 12 | 10.4 | 9.1 | 8.7 | 8.8 |
| LF | *Lactobacillus fermentum* | 12 | 18.4 | 16.1 | 21.6 | 21.9 |
| EC | *Enterococcus faecalis* | 12 | 9.9 | 8.7 | 14.6 | 14.6 |
| SE | *Staphylococcus aureus* | 12 | 15.5 | 13.6 | 15.2 | 15.3 |
| BS | *Listeria monocytogenes* | 12 | 14.1 | 12.4 | 13.9 | 13.9 |
| SC | *Bacillus subtilis* | 12 | 17.4 | 15.3 | 10.3 | 10.3 |
| PA | *Saccharomyces cerevisiae* | 2 | NA | 9.3 | 0.57 | 0.29 |
| LM | *Cryptococcus neoformans* | 2 | NA | 3.3 | 0.37 | 0.18 |

**Table S3**. Composition of ZymoBIOMICS Gut Microbiome Standard.

| Abbreviation | Taxa | Theoretical Composition (%) | | | | |
| --- | --- | --- | --- | --- | --- | --- |
|  |  | Genomic  DNA | 16S Only | 16S & 18S | Genome Copy | Cell Number |
| AM | *Akkermansia muciniphila* | 1.5 | 0.97 | 0.87 | 1.62 | 1.62 |
| BF | *Bacteroides fragilis* | 14 | 9.94 | 9 | 8.33 | 8.36 |
| BA | *Bifidobacterium adolescentis* | 6 | 8.78 | 7.95 | 8.83 | 8.86 |
| CA | *Candida albicans* | 1.5 | N/A | 3.11 | 0.31 | 0.16 |
| CD | *Clostridioides difficile* | 1.5 | 2.62 | 2.37 | 1.1 | 1.1 |
| CP | *Clostridium perfringens* | 0.0001 | 0.0002 | 0.0002 | 0.00009 | 0.00009 |
| EF | *Enterococcus faecalis* | 0.001 | 0.0009 | 0.0008 | 0.0011 | 0.0011 |
| ECB1109 | *Escherichia coli (B-1109)* | 2.8 | 2.46 | 2.23 | 1.77 | 1.77 |
| ECB2207 | *Escherichia coli (B-2207)* | 2.8 | 2.29 | 2.07 | 1.64 | 1.65 |
| ECB3008 | *Escherichia coli (B-3008)* | 2.8 | 2.53 | 2.29 | 1.82 | 1.82 |
| ECB766 | *Escherichia coli (B-766)* | 2.8 | 2.31 | 2.09 | 1.66 | 1.66 |
| ECJM109 | *Escherichia coli (JM109)* | 2.8 | 2.53 | 2.29 | 1.82 | 1.83 |
| FP | *Faecalibacterium prausnitzii* | 14 | 17.63 | 15.96 | 14.77 | 14.82 |
| FN | *Fusobacterium nucleatum* | 6 | 7.49 | 6.79 | 7.53 | 7.56 |
| LF | *Lactobacillus fermentum* | 6 | 9.63 | 8.72 | 9.68 | 9.71 |
| MS | *Methanobrevibacter smithii* | 0.1 | 0.066 | 0.06 | 0.17 | 0.17 |
| PC | *Prevotella corporis* | 6 | 4.98 | 4.51 | 6.26 | 6.28 |
| RH | *Roseburia hominis* | 14 | 9.89 | 8.95 | 12.43 | 12.47 |
| SC | *Saccharomyces cerevisiae* | 1.4 | N/A | 6.35 | 0.32 | 0.16 |
| SE | *Salmonella enterica* | 0.01 | 0.009 | 0.008 | 0.007 | 0.0065 |
| VR | *Veillonella rogosae* | 14 | 15.87 | 14.37 | 19.94 | 20.01 |

**Table S4**. Information of Primers and gblocks for Digital PCR.

| **Target Organism** | **Amplicon length (bp)** | **gene** | **Primer Probe** | **Name** | **Sequence (5'-3')** | **Dyes** | **gBlock standard** | **Anhealing (℃)** | **Threshold (RFU)** |
| --- | --- | --- | --- | --- | --- | --- | --- | --- | --- |
| *Mycobacterium spp.* | 477 | 16S rRNA | forward | I571-110F | CCTGGGAAACTGGGTCTAAT | FAM | A | 55^11^ | 33 |
|  |  |  | reverse | I571R | CGCACGCTCACAGTTA |  |  |  |  |
|  |  |  | probe | Probe H19R | TTTCACGAACAACGCGACAAACT |  |  |  |  |
| *Accumulibacter spp.* | 351 | 16S rRNA | forward | 518f | CCAGCAGCCGCGGTAAT | / | A | 65^12^ | 136 |
|  |  |  | reverse | 846r | GTTAGCTACGGCACTAAAAGG | / |  |  |  |
| *Nitrospira spp.* | 72 | 16S rRNA | forward | Nspra675f | GCGGTGAAATGCGTAGAKATCG | / | B | 60^13^ | 94 |
|  |  |  | reverse | Nspra675r | TCAGCGTCAGRWAYGTTCCAGAG | / |  |  |  |
| *Gordonia spp.* | 829 | 16S rRNA | forward | G268F | CGACCTGAGAGGGTGATCG | / | B | 66^14^ | 73 |
|  |  |  | reverse | G1096r | ATAACCCGCTGGCAATACAG | / |  |  |  |
| A: GGGATAAGCCTGGGAAACTGGGTCTAATACCGAATAGGACCGCGGACTTCATGGTGGTTGTGGTCTCGTAGGTAGTTTGTCGCGTTGTTCGTGAAATCTCACGGCTTAACTGTGAGCGTGCGGGTTTTTCCAGCAGCCGCGGTAATTCCACGTGTAGCGGTGAAATGCGTAGAGATGTGGAGGAACACCGATGGCGAAGGCAGCCCCCTGGGCCAATACTGACGCTCCTTTTAGTGCCGTAGCTAACGCGTGAAG | | | | | | | | | |
| B: GGTGTAGCGGTGAAATGCGTAGAGATCGGGAGGAAGGCCGGTGGCGAAGGCGGCGCTCTGGAACATACCTGACGCTGAGATTTTTGCCGACCTGAGAGGGTGATCGGCCACACTGGGACTGAGACACGGCCCAGACTCCTATGTCCTGTATTGCCAGCGGGTTATGCCGTAT | | | | | | | | | |

**Table S5**. Comparison of relative abundance, mapped bases, and coverage fraction of each genome retrieved using Nanopore and Illumina sequencing platforms under the same sequencing efforts.

|  | Nanopore | | |  | Illumina | | |
| --- | --- | --- | --- | --- | --- | --- | --- |
| Genome | Relative Abundance (%) | mapped bases | Coverage fraction |  | Relative Abundance (%) | mapped bases | Coverage fraction |
| unmapped | 1.14 |  |  |  | 0.15 |  |  |
| *Akkermansia muciniphila* | 2.08 | 1.80E+07 | 99.65% |  | 2.26 | 2.02E+07 | 99.64% |
| *Bacteroides fragilis* | 12.68 | 1.98E+08 | 100.00% |  | 11.22 | 1.82E+08 | 99.89% |
| *Bifidobacterium adolescentis* | 6.07 | 3.84E+07 | 100.00% |  | 8.54 | 5.59E+07 | 100.00% |
| *Candida albican* | 0.20 | 7.48E+06 | 40.96% |  | 0.14 | 5.59E+06 | 30.77% |
| *Clostridioides difficile* | 1.23 | 1.57E+07 | 97.52% |  | 1.09 | 1.44E+07 | 95.27% |
| *Clostridium perfringens* | 0.00 | 0.00E+00 | 0.00% |  | 0.00 | 3.25E+02 | 0.01% |
| *Enterococcus faecalis* | 0.00 | 9.75E+03 | 0.34% |  | 0.00 | 9.09E+03 | 0.30% |
| *Escherichia coli B1109* | 2.65 | 3.91E+07 | 99.42% |  | 2.38 | 3.63E+07 | 99.87% |
| *Escherichia coli b2207* | 2.15 | 3.41E+07 | 99.61% |  | 2.20 | 3.60E+07 | 99.20% |
| *Escherichia coli B3008* | 2.76 | 3.97E+07 | 99.99% |  | 2.41 | 3.58E+07 | 99.86% |
| *Escherichia coli B766* | 2.34 | 3.68E+07 | 97.66% |  | 2.26 | 3.67E+07 | 98.73% |
| *Escherichia coli JM109* | 2.46 | 3.52E+07 | 99.88% |  | 2.33 | 3.45E+07 | 99.81% |
| *Faecalibacterium prausnitzii* | 9.46 | 8.34E+07 | 100.00% |  | 13.86 | 1.26E+08 | 99.98% |
| *Fusobacterium nucleatum* | 6.04 | 4.48E+07 | 100.00% |  | 5.61 | 4.30E+07 | 99.98% |
| *Lactobacillus fermentum* | 5.28 | 3.05E+07 | 100.00% |  | 5.38 | 3.21E+07 | 100.00% |
| *Methanobrevibacter smithii* | 0.22 | 1.26E+06 | 48.17% |  | 0.17 | 9.85E+05 | 38.98% |
| *Prevotella corporis* | 9.63 | 8.60E+07 | 99.98% |  | 8.53 | 7.87E+07 | 99.52% |
| *Roseburia hominis* | 6.89 | 7.22E+07 | 100.00% |  | 9.99 | 1.08E+08 | 99.95% |
| *Saccharomyces cerevisiae* | 0.22 | 8.60E+06 | 46.51% |  | 0.17 | 6.75E+06 | 37.19% |
| *Salmonella enterica* | 0.02 | 2.42E+05 | 4.82% |  | 0.02 | 2.52E+05 | 3.70% |
| *Veillonella rogosae* | 26.48 | 1.73E+08 | 100.00% |  | 21.31 | 1.44E+08 | 99.99% |

**Table S6.** Theoretical relative abundance and coefficient of variation of spike-in controls and ZG samples.

|  | Spike-in controls (ZYMO Log) | | Samples (ZYMO even) | |
| --- | --- | --- | --- | --- |
| Taxa | Theoretical relative abundance (%) | Coefficient of variation (%) | Theoretical relative abundance (%) | Coefficient of variation (%) |
| *Bacillus subtilis* | 0.89 | 4.97 | 10.3 | 0.44 |
| *Cryptococcus neoformans* | 0.00089 | 9.05 | 0.37 | 1.95 |
| *Enterococcus faecalis* | 0.00089 | 3.45 | 14.6 | 3.49 |
| *Escherichia coli* | 0.089 | 4.44 | 8.5 | 2.04 |
| *Lactobacillus fermentum* | 0.0089 | 17.07 | 21.6 | 2.74 |
| *Listeria monocytogenes* | 89.1 | 0.53 | 13.9 | 2.77 |
| *Pseudomonas aeruginosa* | 8.9 | 7.49 | 6.1 | 3.71 |
| *Saccharomyces cerevisiae* | 0.89 | 2.92 | 0.57 | 1.04 |
| *Salmonella enterica* | 0.089 | 2.27 | 8.7 | 1.60 |
| *Staphylococcus aureus* | 0.000089 | 3.99 | 15.2 | 6.64 |

**Table S7**: Mean and median deviations of estimated genome copy number for ZG samples relative to the theoretical values for single spike-in control strategy using different taxa in spike-in controls.

| Type of spike-in controls | Mean deviation | Median deviation | *p_adj* |
| --- | --- | --- | --- |
| Zymo Log | 0.86 | 0.81 |  |
| BF | 0.55 | 0.53 | 1.00E+00 |
| RH | 3.43 | 3.72 | 3.87E-03 |
| EF | 6.19 | 6.53 | 7.59E-02 |
| LF | 8.01 | 9.52 | 2.13E-03 |
| SE | 16.06 | 17.56 | 1.18E-04 |
| CA | 18.57 | 22.55 | 4.02E-07 |
| AM | 0.63 | 0.79 | 1.00E+00 |
| CP | 39549.95 | 7859.15 | 2.15E-12 |
| SC | 10.77 | 12.88 | 4.26E-03 |
| PC | 0.55 | 0.51 | 1.00E+00 |
| FN | 0.56 | 0.54 | 1.00E+00 |
| MS | 6.65 | 6.99 | 5.56E-02 |
| VR | 0.55 | 0.52 | 1.00E+00 |
| FP | 0.93 | 0.89 | 4.65E-01 |
| BA | 10.75 | 13.04 | 4.26E-03 |
| CD | 0.63 | 0.63 | 1.00E+00 |

**Table S8**. Abbreviation for single spike-ins.

| Abbreviation | Taxa | Abbreviation | Taxa | Abbreviation | Taxa |
| --- | --- | --- | --- | --- | --- |
| AM | *Akkermansia muciniphila* | FN | *Fusobacterium nucleatum* | SE | *Salmonella enterica* |
| BA | *Bifidobacterium adolescentis* | FP | *Faecalibacterium prausnitzii* | VR | *Veillonella rogosae* |
| BF | *Bacteroides fragilis* | LF | *Lactobacillus fermentum* |  |  |
| BS | *Bacillus subtilis* | LM | *Listeria monocytogenes* |  |  |
| CA | *Candida albicans* | MS | *Methanobrevibacter smithii* |  |  |
| CD | *Clostridioides difficile* | PA | *Pseudomonas aeruginosa* |  |  |
| CN | *Cryptococcus neoformans* | PC | *Prevotella corporis* |  |  |
| CP | *Clostridium perfringens* | RH | *Roseburia hominis* |  |  |
| EC | *Escherichia coli* | SA | *Staphylococcus aureus* |  |  |
| EF | *Enterococcus faecalis* | SC | *Saccharomyces cerevisiae* |  |  |

**References:**

1.	Chen, S. Ultrafast one-pass FASTQ data preprocessing, quality control, and deduplication using fastp. *iMeta* **2**, e107 (2023).

2.	Danecek, P. *et al.* Twelve years of SAMtools and BCFtools. *GigaScience* **10**, giab008 (2021).

3.	Li, H. Aligning sequence reads, clone sequences and assembly contigs with BWA-MEM. Preprint at https://doi.org/10.48550/arXiv.1303.3997 (2013).

4.	Quinlan, A. R. & Hall, I. M. BEDTools: a flexible suite of utilities for comparing genomic features. *Bioinformatics* **26**, 841–842 (2010).

5.	Vasimuddin, Md., Misra, S., Li, H. & Aluru, S. Efficient Architecture-Aware Acceleration of BWA-MEM for Multicore Systems. in *2019 IEEE International Parallel and Distributed Processing Symposium (IPDPS)* 314–324 (2019). doi:10.1109/IPDPS.2019.00041.

6.	Nurk, S., Meleshko, D., Korobeynikov, A. & Pevzner, P. A. metaSPAdes: a new versatile metagenomic assembler. *Genome Res.* **27**, 824–834 (2017).

7.	Pan, S., Zhao, X.-M. & Coelho, L. P. SemiBin2: self-supervised contrastive learning leads to better MAGs for short- and long-read sequencing. *Bioinformatics* **39**, i21–i29 (2023).

8.	Nissen, J. N. *et al.* Improved metagenome binning and assembly using deep variational autoencoders. *Nat Biotechnol* **39**, 555–560 (2021).

9.	Kang, D. D. *et al.* MetaBAT 2: an adaptive binning algorithm for robust and efficient genome reconstruction from metagenome assemblies. *PeerJ* **7**, e7359 (2019).

10.	Sieber, C. M. K. *et al.* Recovery of genomes from metagenomes via a dereplication, aggregation and scoring strategy. *Nat Microbiol* **3**, 836–843 (2018).

11.	Radomski, N., Kreitmann, L., McIntosh, F. & Behr, M. A. The Critical Role of DNA Extraction for Detection of Mycobacteria in Tissues. *PLOS ONE* **8**, e78749 (2013).

12.	He, S., Gall, D. L. & McMahon, K. D. “Candidatus Accumulibacter” Population Structure in Enhanced Biological Phosphorus Removal Sludges as Revealed by Polyphosphate Kinase Genes. *Appl Environ Microbiol* **73**, 5865–5874 (2007).

13.	Vilardi, K. J. *et al.* Nitrogen source influences the interactions of comammox bacteria with aerobic nitrifiers. *Microbiology Spectrum* **12**, e03181-23 (2024).

14.	Xu, S., Sun, M., Zhang, C., Surampalli, R. & Hu, Z. Filamentous sludge bulking control by nano zero-valent iron in activated sludge treatment systems. *Environ. Sci.: Processes Impacts* **16**, 2721–2728 (2014).
